# SARS-CoV-2 Beta variant infection elicits potent lineage-specific and cross-reactive antibodies

**DOI:** 10.1101/2021.09.30.462420

**Authors:** S Momsen Reincke, Meng Yuan, Hans-Christian Kornau, Victor M Corman, Scott van Hoof, Elisa Sánchez-Sendin, Melanie Ramberger, Wenli Yu, Yuanzi Hua, Henry Tien, Marie Luisa Schmidt, Tatjana Schwarz, Lara Maria Jeworowski, Sarah E Brandl, Helle Foverskov Rasmussen, Marie A Homeyer, Laura Stöffler, Martin Barner, Désirée Kunkel, Shufan Huo, Johannes Horler, Niels von Wardenburg, Inge Kroidl, Tabea M Eser, Andreas Wieser, Christof Geldmacher, Michael Hoelscher, Hannes Gänzer, Günter Weiss, Dietmar Schmitz, Christian Drosten, Harald Prüss, Ian A. Wilson, Jakob Kreye

**Affiliations:** Charité-Universitätsmedizin Berlin, corporate member of Freie Universität Berlin and Humboldt-Universität zu Berlin, Department of Neurology and Experimental Neurology, Berlin, Germany; German Center for Neurodegenerative Diseases (DZNE) Berlin, Berlin, Germany; Helmholtz Innovation Lab BaoBab (Brain antibody-omics and B-cell Lab), Berlin, Germany; Department of Integrative Structural and Computational Biology, The Scripps Research Institute, La Jolla, CA 92037, USA; Neuroscience Research Center (NWFZ), Cluster NeuroCure, Charité-Universitätsmedizin Berlin, corporate member of Freie Universität Berlin, Humboldt-Universität Berlin, and Berlin Institute of Health, Berlin, Germany; Charité-Universitätsmedizin Berlin, corporate member of Freie Universität Berlin and Humboldt-Universität zu Berlin, Institute of Virology, Berlin, Germany and German Centre for Infection Research (DZIF), Berlin, Germany; Labor Berlin – Charité Vivantes GmbH, Berlin; Berlin Institute of Health at Charité - Universitätsmedizin Berlin, Flow & Mass Cytometry Core Facility, Berlin, Germany; Division of Infectious Diseases and Tropical Medicine, Medical Center of the University of Munich (LMU), Germany; German Center for Infection Research (DZIF), partner site Munich, Germany; Department of Internal Medicine, BKH Schwaz, Schwaz, Austria; Department of Internal Medicine II, Medical University of Innsbruck, Innsbruck, Austria; The Skaggs Institute for Chemical Biology, The Scripps Research Institute, La Jolla, CA 92037, USA; Charité-Universitätsmedizin Berlin, corporate member of Freie Universität Berlin and Humboldt-Universität zu Berlin, Department of Pediatric Neurology, Berlin, Germany

**Author notes:** These authors contributed equally to this work (S Momsen Reincke, Meng Yuan, Hans-Christian Kornau, Victor M Corman). These authors contributed equally to this work (Harald Prüss, Ian A. Wilson, Jakob Kreye). Corresponding authors (S.M.R.); (H.P.); (I.A.W.); (J.K.).

## Abstract

SARS-CoV-2 Beta variant of concern (VOC) resists neutralization by major classes of antibodies from non-VOC COVID-19 patients and vaccinated individuals. Here, serum of Beta variant infected patients revealed reduced cross-neutralization of non-VOC virus. From these patients, we isolated Beta-specific and cross-reactive receptor-binding domain (RBD) antibodies. The Beta-specificity results from recruitment of novel VOC-specific clonotypes and accommodation of VOC-defining amino acids into a major non-VOC antibody class that is normally sensitive to these mutations. The Beta-elicited cross-reactive antibodies share genetic and structural features with non-VOC-elicited antibodies, including a public VH1-58 clonotype targeting the RBD ridge independent of VOC mutations. These findings advance our understanding of the antibody response to SARS-CoV-2 shaped by antigenic drift with implications for design of next-generation vaccines and therapeutics.

**One sentence summary:** SARS-CoV-2 Beta variant elicits lineage-specific antibodies and antibodies with neutralizing breadth against wild-type virus and VOCs.

## MAIN

In the course of the COVID-19 pandemic, multiple SARS-CoV-2 lineages have emerged including lineages defined as variants of concern (VOC), such as Alpha (also known as lineage B.1.1.7), Beta (B.1.351), Gamma (P.1) and Delta (B.1.617.2). VOCs are associated with increased transmissibility, virulence and/or resistance to neutralization by sera from vaccinated individuals and convalescent COVID-19 patients who were infected with the original, non-VOC strain (*1–7*). These distinct lineages carry a variety of mutations in the spike protein, several of which are within the receptor binding domain (RBD), especially at residues K417, L452, T478, E484, and N501. Some mutations like N501Y are associated with enhanced binding to angiotensin-converting enzyme 2 (ACE2), largely driving the global spread of VOCs incorpororating these mutations (*2*). However, with increasing immunity either through natural infection or vaccination, antibody escape might become more relevant in emerging VOCs. Many studies have investigated RBD antibodies in COVID-19 patients prior to identification of SARS-CoV-2 variants, and we refer to these as non-VOC antibodies. Non-VOC RBD antibodies revealed a preferential response towards distinct epitopes with enriched recruitment of particular antibody germline genes, where the most prominent were VH3-53 and closely related VH3-66, as well as VH1-2 (*8, 9*). Structural and functional classification of non-VOC RBD mAbs has demonstrated that mAbs from these three enriched germline genes form two major classes of receptor-binding site (RBS) mAbs whose binding and neutralizing activity depends on either K417 and/or E484 (*9, 10*). Mutations at these key residues (K417N and E484K) and at N501Y are hallmarks of the Beta variant (*2*), and largely account for the reduced neutralizing activity of sera from vaccinated individuals and convalescent COVID-19 patients against this VOC (*1–6, 11, 12*). These key mutations also occur in further sublineages including an Alpha variant strain carrying E484K and a Delta variant strain carrying K417N (Delta Plus). Of all current VOCs, the Beta variant appears to be most resistant to neutralization from non-VOC sera, suggesting conspicuous differences in its antigenicity (*13*). However, little is known about the antibody response elicited by Beta variant infection. For example, it is unknown if antibodies targeting the RBD of the Beta variant (RBD Beta) share the preferential recruitment of particular germline genes with non-VOC antibodies, and whether VOC-defining mutations K417N and E484K could be accommodated in the canonical binding modes of public antibody classes like VH3-53/VH3-66 antibodies. Investigating Beta variant induced immunity can therefore bolster efforts to monitor and prevent the spread of SARS-CoV-2 by informing vaccine design in the context of the ongoing antigenic drift. Thus, we set out to explore genetic, functional and structural features of the antibody response against RBD Beta compared to non-VOC RBD.

We identified 40 individuals infected with the SARS-CoV-2 Beta variant from three metropolitan areas in Germany and Austria (table S1). Serum from these patients was collected 38.6 ± 19.2 days after their first positive SARS-CoV-2 RT-PCR test. The patients’ IgG bound to non-VOC nucleocapsid protein and/or non-VOC spike protein in 37 of 40 patients with stronger reactivity to RBD Beta than to non-VOC RBD (fig. S1A). The VOC patients’ sera also inhibited ACE2 binding to RBD Beta to a greater extent than to non-VOC RBD (fig. S1B, table S1), indicating the presence of highly effective RBD antibodies after Beta variant infection. Reactivity to non-VOC spike S1 antigen was confirmed in an additional commercially available ELISA; however, only 23 of 40 samples tested positive according to the manufacturer’s cutoff (fig. S1, C and D). In a plaque reduction neutralization test (PRNT), 37 of 40 sera neutralized an authentic SARS-CoV-2 Beta isolate (B.1.351) with a half-maximal inhibitory concentration (IC_50_) at 1:20 dilution or greater (Fig. 1A). On the other hand, only 11 of 40 sera neutralized non-VOC authentic virus of Munich isolate 984 (*14*) with an IC_50_ at 1:20 dilution or greater (Fig. 1B). The neutralizing activity against the two viral isolates was modestly correlated (fig. S1E), with a ~20.4-fold reduction of neutralizing activity against the non-VOC virus compared to Beta as measured by the area under the curve (AUC) of neutralization titers (Fig. 1C), indicating limited breadth of serum antibodies in our cohort. A converse effect has been reported after immune responses against non-VOC RBD in cohorts of convalescent patients and of vaccinated individuals (*2*), where neutralization of SARS-CoV-2 Beta was ~8 to ~14 fold reduced compared to non-VOC virus (*1–6*). No positive correlation was found between neutralizing antibodies against Beta and the timepoint of sample collection relative to first positive PCR test (fig. S1F). Neutralizing antibodies against Beta modestly correlated with age (fig. S1G), but no statistically significant gender difference was observed (fig. S1H). Collectively, these data show that sera from Beta-infected patients exhibit reduced cross-reactivity and cross-neutralization to non-VOC SARS-CoV-2, impacting diagnostic antibody testing and assessment of antibody levels when using non-VOC antigens.

**Fig. 1.**
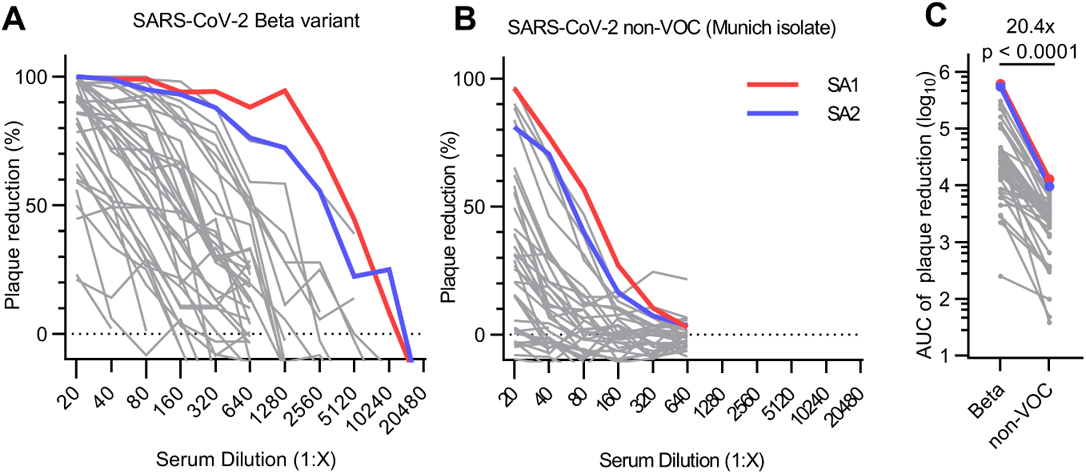
Authentic virus neutralization of sera from individuals after infection with SARS-CoV-2 Beta. **(A-B)** Neutralizing activity of sera of patients infected with SARS-CoV-2 Beta variant was measured using a plaque-reduction neutralization assay with the indicated authentic virus. Results are given as reduction of plaque number at indicated serum dilutions. Patient SA1 and SA2 are highlighted in red and blue, respectively. Means of duplicate measurements are shown. Values below zero indicate no plaque reduction. **(C)** Change in neutralization activity against SARS-CoV-2 Beta and non-VOC SARS-CoV-2 based on area under the curve (AUC) calculations from authentic virus PRNT curves (shown in **A** and **B**). Mean fold change is indicated above the p value. Statistical analysis was performed using a Wilcoxon matched-pairs signed-rank test with two-tailed p value.

To investigate this difference in reactivity between RBD Beta and non-VOC RBD on the level of mAbs elicited by SARS-CoV-2 Beta variant infection, we isolated CD19^+^CD27^+^ memory B cells from the peripheral blood of 12 donors in our cohort via fluorescence activated cell sorting using a recombinant RBD Beta (K417N/E484K/N501Y) probe (fig. S2A). Frequencies of RBD-double-positive memory B cells ranged from 0.007% to 0.1% (fig. S2B). Using single-cell Ig gene sequencing (*15, 16*), we derived 289 pairs of functional heavy (IGH) and light (IGL) chain sequences from IgG mAbs (table S2). Sequence analysis showed enrichment of certain VH genes compared to mAbs derived from healthy, non-infected individuals, including VH1-58, VH3-30, VH4-39 and VH3-53, illustrating a preferential recruitment of certain VH genes (Fig. 2A) and VH-JH gene combinations (fig. S3A). For some genes like VH1-58 and VH3-53, enrichment has previously been identified in CoV-AbDab, a database of published non-VOC SARS-CoV-2 mAbs (*9, 17*). We here confirmed this finding for all human RBD mAbs in this database (Fig. 2A). Consistent with previous reports from non-VOC SARS-CoV-2 infections (*18–20*), the number of somatic hypermutations (SHM) in the IGH and IGL chains was generally low in mAbs derived from our cohort (fig. S3B). Together, these findings argue for conservation of certain antibody sequence features between antibody responses in different donors and between antibody responses elicited against SARS-CoV-2 Beta variant and non-VOC viruses. Hence, we compared antibody sequences after Beta infection to all previously published non-VOC RBD antibodies and identified several clonotypes shared between both datasets (Fig. 2B), some of which were present in multiple patients of our study (Fig. 2C). Taken together, these results demonstrate that a subset of the antibodies to RBD Beta and non-VOC RBD converge on recruitment of specific germline genes.

**Fig. 2.**
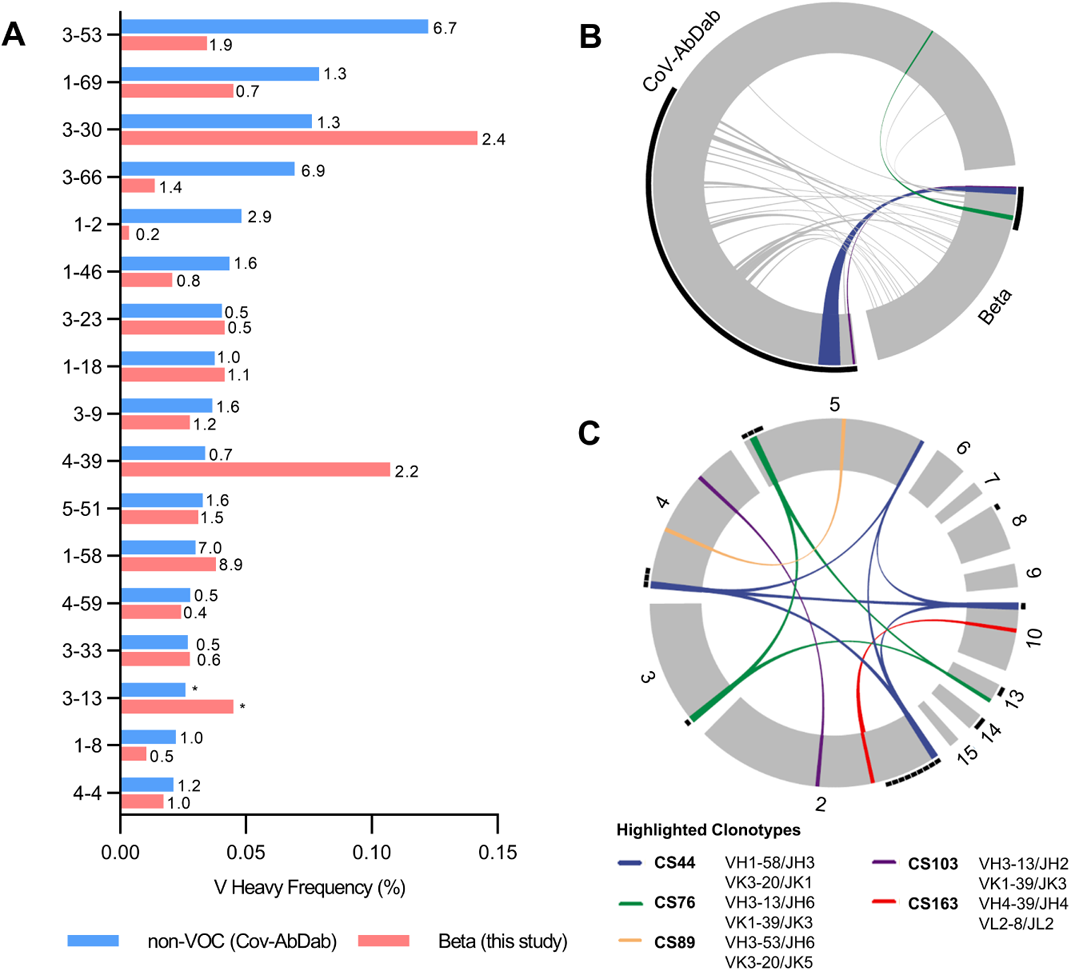
Germline gene usage and clonotype analysis of Beta-elicited antibodies. **(A)** VH gene usage of 289 RBD Beta IgG mAbs from this study (red) is compared to 1037 non-VOC RBD mAbs from 96 previously published studies (blue, Cov-AbDab) (*17*). Frequencies of mAbs encoded by each VH gene are shown as bars. Enrichment of indicated VH genes is compared to healthy individuals (*31*) with fold-enrichment shown as number next to bars. VH gene frequencies that were not reported in healthy individuals (*31*) are shown with asterisk (*). Only VH genes with a frequency of at least 2% in Cov-AbDab are shown and VH genes are ordered by frequency in Cov-AbDab. **(B**) Circos plot shows the relationship between 289 IgG mAbs from this study (Beta) and 1037 previously published human mAbs reactive to RBD (non-VOC) from 96 studies (*17*). Interconnecting lines display clonotypes shared between both datasets, as defined by the usage of the same V and J gene on both Ig heavy and light chain. Thin black lines at the outer circle border indicate expanded clonotypes within the respective data set. **(C)** Circos plot displaying the 289 IgG mAbs from this study grouped per patient. Interconnecting colored lines indicate clonotypes found in more than one patient. Small black angles at the outer circle border indicate clonally expanded clones within one patient. **(B-C)** Colored interconnecting lines depict clonotypes found in more than one patient of our cohort.

However, other gene enrichments found in our study like VH4-39 have not been identified within the CoV-AbDab mAbs (*9*) (Fig. 2A), exemplifying concurrent divergence in the antibody response to the different RBDs. Strikingly, VH1-2, one of the most common genes contributing to the RBD antibody response to non-VOC SARS-CoV-2, was virtually absent in our dataset of Beta variant elicited mAbs (Fig. 2A and table S2), in line with our previous predictions about the effect of Beta variant mutations on VH1-2 binding and neutralization (*9*). VH3-53/VH3-66 antibodies have previously been shown to bind to non-VOC RBD in two canonical binding modes, which involve residues K417 and E484, respectively; binding and neutralization of these antibodies are strongly affected by the K417N and E484K mutations in RBD Beta (*9, 21*). We therefore hypothesized a similarly reduced recruitment of VH3-53/VH3-66 mAbs after Beta variant infection. Surprisingly, we identified 15 VH3-53/VH3-66 mAbs, albeit at a reduced frequency compared to the CoV-AbDab dataset (4.7% vs. 19.4%), but still at an increased frequency compared to healthy donors (Fig. 2A), thus indicating either a non-canonical binding mode or accommodation of these mutations into the known binding modes.

To determine the binding properties of antibodies elicited by SARS-CoV-2 Beta, we selected mAbs for expression and further characterization based on the following criteria: (i) mAbs that are clonally expanded within one patient, (ii) mAbs of clonotypes present in several patients in our dataset to decipher the shared antibody response to RBD Beta, (iii) mAbs of clonotypes found both in our and CoV-AbDab datasets to potentially identify cross-reactive mAbs, (iv) VH3-53/VH3-66 mAbs to elucidate the unexpected recurrent shared antibody response against RBD Beta, (v) mAbs of VH genes with strongest enrichment in our dataset, including VH4-39 and VH1-58. We identified 81 mAbs with strong binding to RBD Beta (table S3), as defined by detectable binding at 10 ng/ml. Of those, a majority (44 in total) revealed comparable binding to non-VOC RBD and were considered cross-reactive mAbs, whereas 37 mAbs did not bind non-VOC RBD at 10 ng/ml and were considered RBD Beta-specific.

We aimed to determine the residues that define the mAb binding selectivity for the 37 RBD Beta-specific mAbs, and performed ELISAs with single mutant constructs of RBD Beta and non-VOC RBD. For all three Beta-defining RBD mutations (K417N, E484K and N501Y), we identified mAbs with RBD binding that were dependent on a single residue, with a larger fraction of binders that were dependent on K484 (12) and Y501 (11) than N417 (3). The RBD Beta specificity of other mAbs was dependent on multiple residues (Fig. 3A). 26 of the RBD Beta-specific mAbs (70.2%) neutralized the authentic SARS-CoV-2 Beta isolate, with representation in all of the above-mentioned specificity categories (Fig. 3A). RBD Beta-specific mAbs were encoded by a broad variety of VH genes (Fig. 3A and table S2). Interestingly, all nine VH4-39 mAbs with RBD Beta specificity from three different patients were Y501-dependent, comprising 81.8% of the mAbs in this category. This finding suggests a common binding mode of these mAbs that depends on Y501, a mutation that is present in RBD Beta, Alpha and Gamma, but not Delta, and may explain the frequent use of VH4-39 in mAbs to RBD Beta (Fig. 2A). VH4-39 Y501-dependent mAbs revealed few SHM in VH genes but no uniform pattern in other sequence features (fig. S4A). Although all VH4-39 RBD Beta-specific mAbs bind to a Y501-dependent epitope, their neutralization activity against authentic Beta virus showed noticeable differences (IC_50_ ranging from 5.2 to 947 ng/ml, fig. S4B). Surface plasmon resonance measurements of these mAbs to RBD Beta revealed equilibrium dissociation constants (K_D_) between 3.39 and 80.4 nM (fig. S5A) with correlation to their PRNT-derived IC_50_ values (fig. S5B), thereby providing an explanation for the variability in neutralizing activity within the VH4-39 Y501-dependent mAbs.

**Fig. 3.**
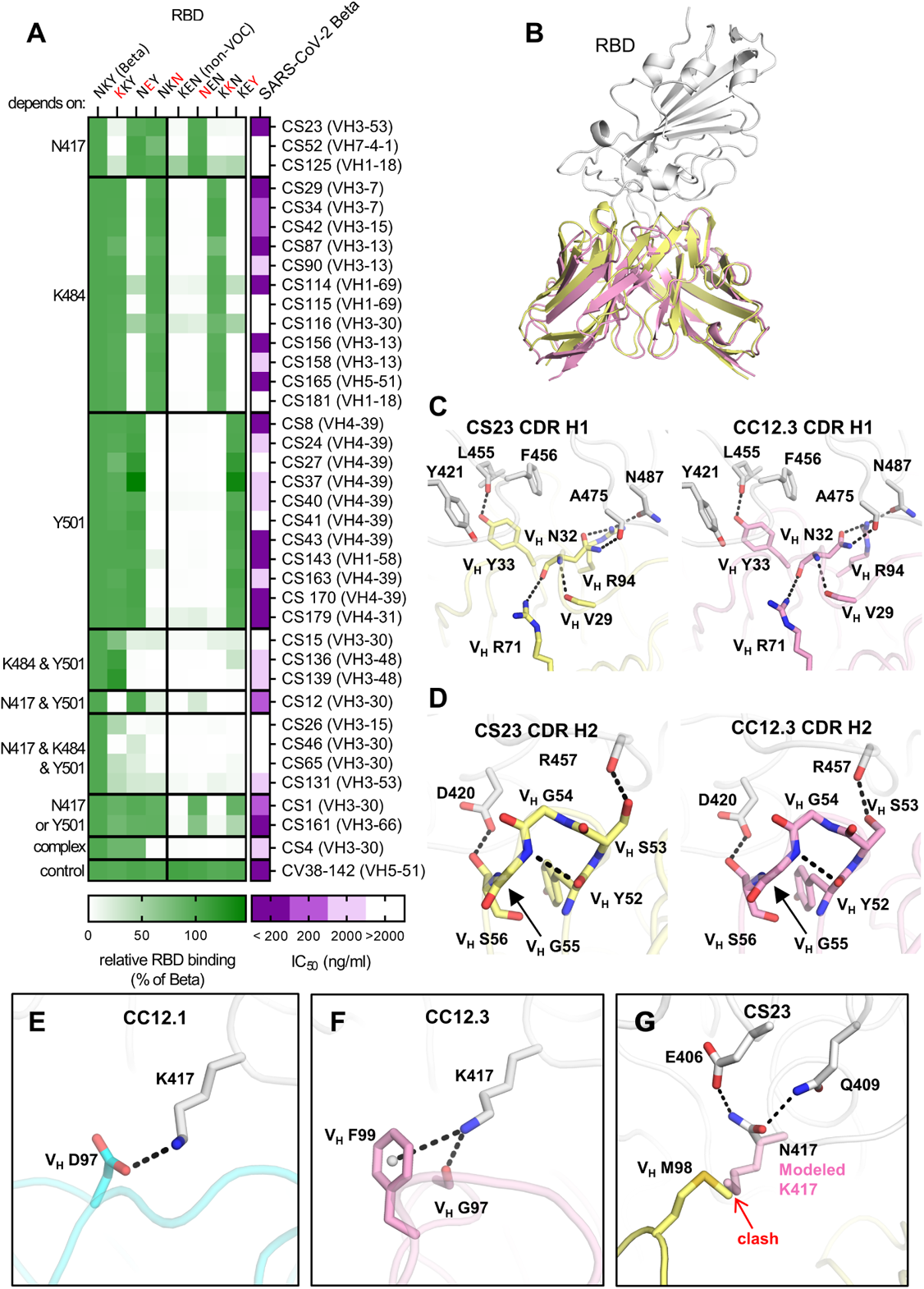
Binding, neutralization and structures of Beta-specific antibodies. **(A)** Neutralization of indicated Beta-specific mAbs against authentic Beta virus is shown in purple. Binding to single point mutant RBD constructs with the indicated amino-acid residues at positions 417, 484 and 501 is shown in green, normalized to RBD Beta. **(B-G)** Structural comparison of VH3-53 mAbs between Beta-specific CS23 and non-VOC-specific CC12.1 and CC12.3. **(B)** CC12.3 and CS23 adopt the same binding mode. The crystal structure of CC12.3 (pink) in complex with RBD (non-VOC) was superimposed onto CS23 (yellow) in complex with RBD (Beta). Only the variable domains of the antibodies are shown for clarity. A small local conformational difference was observed between CS23-bound Beta-RBD and CC12.3-bound non-VOC-RBD (191 Cα, RMSD = 0.8 Å). **(C-D)** Comparison of the **(C)** CDR H1 (‘NY’ motif) and **(D)** CDR H2 (‘SGGS’ motif) between CS23 and CC12.3. **(E-G)** Structures of CDR H3 of **(E)** CC12.1, **(F)** CC12.3, and **(G)** CS23. A modeled side chain of K417 is shown as transparent pink sticks, which would be unfavorable for binding to CS23, where V_H_ M98 occupies this pocket. Structures of CC12.1 (PDB 6XC3, cyan), CC12.3 (PDB 6XC4, pink), and CS23 (this study, yellow) are used throughout this figure, and the RBD is shown in white. Hydrogen bonds, salt bridges or cation-π bonds are represented by black dashed lines.

Furthermore, we identified three VH3-53/VH3-66 mAbs with RBD Beta specificity that all showed neutralizing activity. To determine whether this RBD Beta specificity results from a non-canonical binding mode or accommodation of the Beta variant-defining mutations in one of the two main VH3-53/VH3-66 mAb binding modes, we determined a crystal structure of VH3-53 antibody CS23 in complex with RBD Beta. Previously, we and others found that VH3-53/VH3-66 mAbs with short CDRs H3 (<15 amino acids) target the RBS of non-VOC RBD via a canonical mode (*10, 22–25*) that is highly sensitive to the K417N mutation (*9*). CS23 contains a short CDR H3 with only ten amino acids and is specific to N417 RBDs including RBD Beta (Fig. 3A). Perhaps unexpectedly, CS23 binds to RBD Beta in the canonical mode, with a nearly identical approach angle compared to non-VOC VH3-53 antibody CC12.3 (*24*) (Fig. 3B). We also previously determined that the CDR H1 ^33^NY^34^ and H2 ^53^SGGS^56^ motifs of VH3-53/VH3-66 mAbs are critical for RBD recognition (*24*). Here we find that CS23 retains these motifs and they interact with the RBD in the same way (Fig. 3, C and D). Residues in CDR H3 usually interact with K417 and thus confer specificity to the non-VOC RBD (*9*). For example, V_H_ D97 of CC12.1 forms a salt bridge with the outward facing RBD-K417, whereas V_H_ F99 and V_H_ G97 of CC12.3 interact with K417 through cation-π and hydrogen bonds (H-bonds), respectively (Fig. 3, E and F). Instead, in RBD Beta, the shorter N417 flips inward and H-bonds with RBD-E406 and Q409 (Fig. 4G). V_H_ M98 occupies the vacated space and interacts with RBD-Y453, L455, and V_L_ W91 in a hydrophobic pocket. We modeled in K417 and found that it would be unfavorable for RBD binding to CS23, as V_H_ M98 now occupies the pocket that K417 normally accesses. CDR H3 contains a V_H_ ^96^TAMA^99^ sequence that forms an ST motif that stabilizes CDR H3 and the disposition of M98. The first serine (S) or threonine (T) residue in a four of five residue ST motif makes two internal H-bonds from the side-chain oxygen of residue i to the main-chain NH of residue i + 2 or i + 3, and between the main-chain oxygen of residue i and the main-chain NH of residue i + 3 or i + 4. In this case, the T96 side chain H-bonds with the main-chain NH of M98, and the T96 main-chain oxygen H-bonds with the main-chain NH of A99 (fig. S6A). In fact, this V_H_ ^96^TxMx^99^ motif is unique in all CDRs H3 of anti-RBD VH3-53/VH3-66 antibodies (*17*) and explains the newly acquired specificity of this VH3-53 antibody for an RBD with N417. Previously, non-VOC VH3-53 antibody COVOX-222 was shown to cross-react with RBD Beta but still bind the RBD in the canonical mode; in this case, a rare SHM V_L_ S30P mutation accommodated Y501 (*1*) (fig. S6, B and C). Likewise, for CS23, the CDR L1 ^30^SK^31^ dipeptide is mutated to ^30^GQ^31^ and accommodates Y501 in the Beta variant (fig. S6D). Interestingly, another VH3-53 antibody of our cohort, CS82 is highly cross-reactive and resistant to all tested VOCs, including those with mutations at residues 417 and 484 (Fig. 4A). Moreover, CS82 also binds SARS-CoV (Fig. 4A), which is unprecedented for any previous VH3-53 antibodies. CS82 competes with CR3022 (fig. S6E), suggesting a possible alternative binding interaction compared to the two binding modes of non-VOC VH3-53 antibodies that are sensitive to K417N and E484K, respectively (*23, 24*). Collectively, VH3-53/VH3-66 mAbs contribute to the immune response to RBD Beta with mAbs that accommodate Beta-specific mutations into canonical modes and by mAbs that may bind in alternative binding modes.

**Fig. 4.**
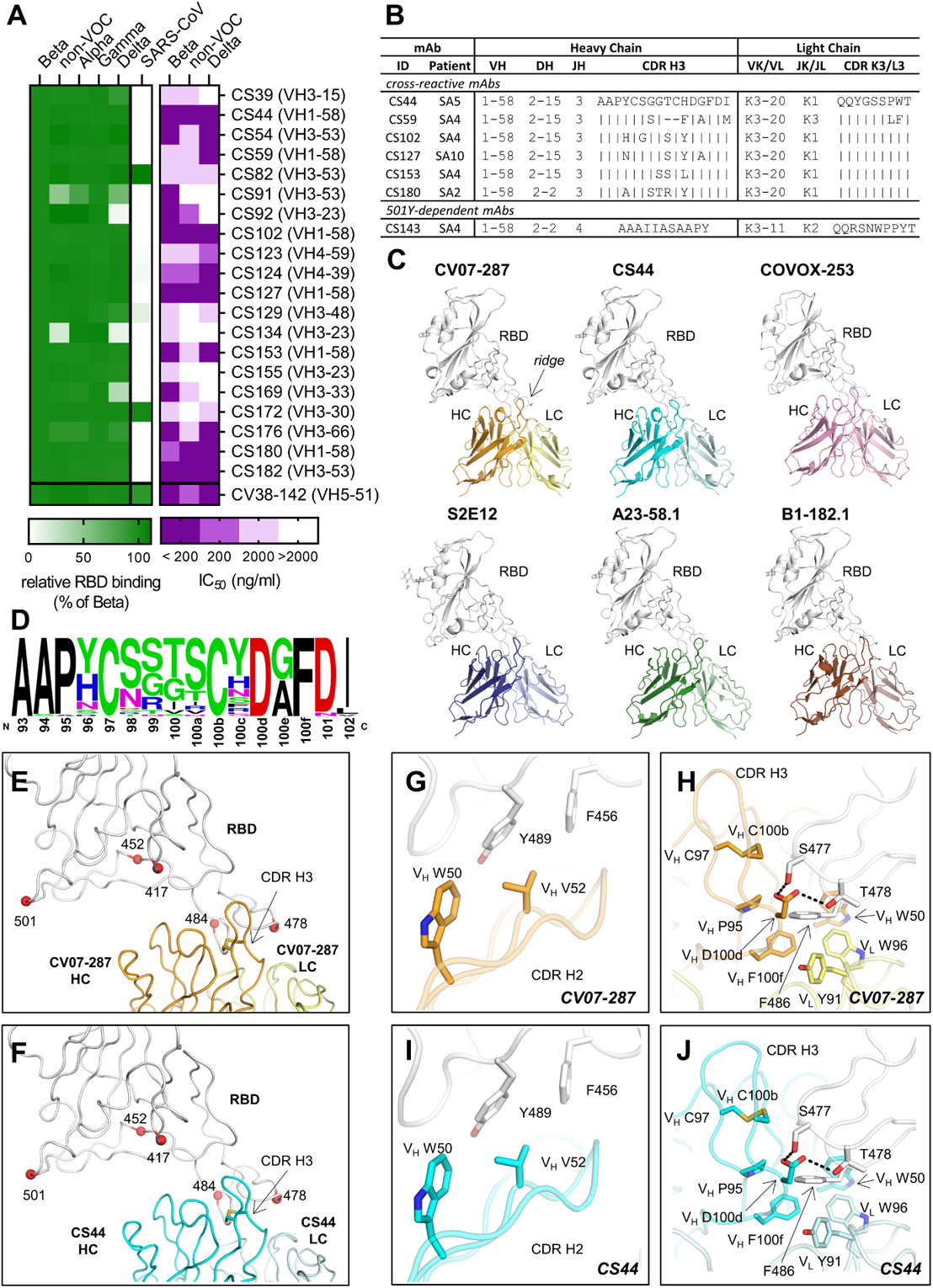
Characterization of cross-reactive mAbs and crystal structures of CV07-287 and CS44. **(A)** Neutralization of cross-reactive antibodies against authentic Beta, Delta and non-VOC virus is shown in purple. Binding to the indicated RBD constructs is shown in green, normalized to RBD Beta. **(B)** Comparison of sequence features from VH1-58 mAbs. **(C)** VH1-58 antibodies target SARS-CoV-2 RBD via the same binding mode. Crystal structures of CV07-287 in complex with non-VOC RBD and CS44 in complex with RBD Beta are shown. COVA1-16 Fab that was used in the crystallization to form the crystal lattice is not shown for clarity. Structures of VH1-58 antibodies from other studies are shown for comparison, including COVOX-253 (PDB 7BEN), S2E12 (PDB 7K45), A23-58.1 (PDB 7LRT), and B1-182.1 (PDB 7MM0). All structures are shown in the same orientation, with the constant domains of the Fab omitted for clarity. The location of the ridge region of the RBD is indicated in the first panel. of the Fab omitted for clarity. The location of the ridge region of the RBD is indicated in the first panel. **(D)** Sequence logo of CDR H3 of VH1-58/VK3-20 antibodies. CDR H3 sequences of VH1-58/VK3-20 antibodies from COVID-19 patients (*17*) were aligned and analyzed with WebLogo. **(E-F)** Mutated residues in VOCs B.1.1.7 (Alpha), B.1.351 (Beta), B.1.617.2 (Delta), and P.1 (Gamma) variants are represented by red spheres. All of these residues are distant from VH1-58 antibodies **(E)** CV07-287 and **(F)** CS44, except for T478. The disulfide bond in each CDR H3 is shown as sticks. **(G-J)** Detailed interactions between the RBD and **(G-H)** CV07-287, and **(I-J)** CS44, respectively. RBDs are shown in white, with heavy and light chains of CV07-287 in orange and yellow, and those of CS44 in cyan and light cyan, respectively. Interactions of CDR H2 are shown in panels **(G)** and **(I)**, and those of CDR H3 are in panels **(H)** and **(J)**. Hydrogen bonds are represented by black dashed lines.

We next aimed to characterize the functional breadth of the cross-reactive mAbs. 20 of the 44 cross-reactive mAbs (45.5%) neutralized authentic SARS-CoV-2 Beta isolate (Fig. 4A). To investigate their cross-reactivity against further RBD variants, we performed ELISAs with RBD constructs of all VOCs and also SARS-CoV. Whereas only two mAbs (10%) strongly detected SARS-CoV RBD, the majority of cross-reactive antibodies bound the RBD of Alpha, Gamma and Delta (Fig. 4A). In PRNT assays with further authentic virus isolates, 15 (75%) Beta variant neutralizing cross-reactive mAbs also neutralized non-VOC virus and 14 (70%) neutralized a Delta virus isolate (Fig. 4A). Thus, cross-reactive mAbs elicited from SARS-CoV-2 Beta infections revealed breadth in binding and neutralization of current VOCs. Six cross-neutralizing antibodies were encoded by VH1-58 (Fig. 4A). VH1-58 is the most enriched germline VH gene in RBD antibodies in both Beta variant and non-VOC infection (*26*) (Fig. 2A). In the context of RBD antibodies, VH1-58 almost exclusively pairs with JH3 (fig. S3A). The VH1-58/JH3/VK3-20/JK1 clonotype has been described in non-VOC infected individuals (*20, 26*) and found in several patients within our cohort (Fig. 2C and Fig. 4B), thereby representing 2.4% of all IgG mAbs analyzed in this study (table S2). To elucidate the structural basis of this public pan-VOC clonotype, we determined crystal structures of CS44 and CV07-287, a mAb of the same clonotype that was isolated from a non-VOC infected individual (*19*), in complex with RBD Beta and non-VOC RBD respectively (Fig. 4C). We compared the structures of CS44 and CV07-287 with other published VH1-58 antibodies, including COVOX-253 (*27*), S2E12 (*28*), A23-58.1, and B1-182.1 (*26*). These antibodies all target the RBD in the same binding mode (Fig. 4C), which suggests that this public clonotype is structurally conserved. The dominant interaction of VH1-58 antibodies is with the RBD ridge region (residues 471–491), which accounts for ~75% of the entire epitope surface. Most of the VOC mutations occur outside of the ridge region (e.g. residues 417, 452, and 501) and are distant from the binding sites of VH1-58 antibodies CV07-287 and CS44 (Fig. 4, E and F). Among the VOC-related residues, only T478 interacts with VH1-58 antibodies, but mutation to a lysine can be accommodated (Fig. 4A) (*26*). V_H_ W50 and Y52 in CDR H2 provide hydrophobic interactions with the RBD (Fig. 4, G and I). CDR H3 also forms extensive interactions with the RBD. The CDR H3 sequences of 38 antibodies that belong to this clonotype (*17*) (Fig. 4D) are highly conserved, and all contain a disulfide bond between V_H_ C97 and C100b, with four relatively small residues (G, S, T) in-between (Fig. 4, D, H and J). V_H_ D100d is also conserved (Fig. 4D), forming H-bonds with S477 and T478 (Fig. 4, H and J). In addition, the conserved V_H_ P95 and F100f (Fig. 4D) stack with RBD-F486 together with V_H_ W50, V_L_ Y91, and V_L_ W96 (Fig. 4, H and J). While E484 is often an important residue for antibody binding on the ridge region, here it is 5 Å distant from the antibodies and mutations at this site have not been reported as being sensitive for VH1-58 antibodies.

Thousands of anti-SARS-CoV-2 mAbs were isolated before the VOCs started to emerge (*17*), many of which are highly potent but sensitive or resistant to VOCs. Here, we characterized the antibody response to the RBD after SARS-CoV-2 Beta variant infection to provide insights into diverging and converging features of antibodies elicited by this lineage compared to non-VOC-elicited antibodies. Our analysis of polyclonal patient sera indicates that non-VOC RBD based diagnostics may underestimate antibody titers after Beta variant infection or even result in a false-negative serological assessment of a prior infection. This issue could be addressed by the use of VOC-based antigens, but these data challenge the concept of defining a universal threshold for protective antibody titers or threshold-based reasoning for booster vaccination.

Furthermore, on the monoclonal level, we show that RBD mAbs with Beta-specificity are frequent after Beta infection, similar to the frequency of RBD mAbs that do not react with RBD Beta after non-VOC infection or vaccination (*2*). Antibodies with Beta-specificity include novel VOC-specific public clonotypes such as Y501-dependent VH4-39 mAbs, and others in the major non-VOC antibody class encoded by VH3-53/VH3-66 genes. Surprisingly, VOC-defining mutations in the Beta variant can be accommodated by local conformational changes and mutations in these VH3-53/VH3-66 mAbs that enable canonical mode binding. Moreover, a subset of RBD Beta elicited mAbs was cross-reactive to non-VOC RBD and also against the other VOCs investigated including the Delta variant, with multiple of these mAbs revealing potent cross-neutralization. VH1-58 antibodies form a clonotype with both ultra-high potency and high resistance to currently circulating VOCs, including variants of concern Alpha, Beta, Gamma and Delta (*26*). Here, we show that pan-VOC VH1-58 antibodies are also frequently elicited by the Beta variant, and target the S protein in a nearly identical binding mode compared to those isolated from non-VOC-infected patients. In fact, VH1-58 is the most enriched germline gene in antibodies isolated from this cohort and may play an important role in neutralizing a broad spectrum of variants. As the epitope of VH1-58 antibodies remains a site of vulnerability in all currently circulating VOCs, these findings provide valuable insights for next-generation vaccine design and antibody therapeutics against present and future variants. For example, simultaneous and/or sequential immunization with vaccines based on diverse RBD sequences could be evaluated for superiority in induction of cross-variant immunity including recruitment of highly potent VH1-58 antibodies. Novel vaccine candidates based on the Beta variant have already showed promising cross-variant antibody titers in pre-clinical studies (*29, 30*), which led to the subsequent initiation of a phase II/III clinical trial. These clinical trials should be complemented by studies characterizing the immune response against further SARS-CoV-2 variants including the Delta variant, which is currently the globally dominating SARS-CoV-2 VOC.

## ACKNOWLEDGEMENTS

We thank all study participants who devoted samples and time to our research, Dr. Kim Stahlberg for patient recruitment, Stefanie Bandura, Matthias Sillmann, Doreen Brandl, Patricia Tscheak, and Sabine Engl for excellent technical assistance, and Dr. Marcel A Müller and Dr. Daniela Niemeyer for support with BSL3 work. We acknowledge BIAFFIN GmbH & Co. KG (Kassel, Germany) for performance of SPR measurements and the Flow & Mass Cytometry Core Facility at Charité-Universitätsmedizin Berlin for support with single-cell sorting. We thank Robyn Stanfield for assistance in data collection, Fangzhu Zhao for assistance in the biolayer interferometry binding assay, and the staff of Advanced Light Source beamline 5.0.1 and Stanford Synchrotron Radiation Laboratory (SSRL) beamline 12-1 for assistance. SARS-CoV-2 RBD variants antigens for sera testing were kindly provided by InVivo BioTech Services GmbH (Hennigsdorf, Germany) to the Seramun Diagnostica GmbH (Heidesee, Germany). S.M.R. and J.K. are participants in the BIH-Charité Junior Clinician Scientist Program and V.M.C is supported by Berlin Institute of Health (BIH) Charité Clinician Scientist program both funded by Charité – Universitätsmedizin Berlin and the Berlin Institute of Health.

## Supplementary Materials

Materials and Methods Figs. S1 to S6

Tables S1 to S5 References 32-44

## Author contributions

Conceptualization: S.M.R., M.Y., H.-C.K., V.M.C., H.P., I.A.W., and J.K.; Patient recruitment and sample preparation: S.M.R., M.R., M.A.H., I.K., T.M.E., C.G., A.W., M.H., H.G., G.W., S.H., H.P., and J.K.; Antibody production: S.M.R., S.v.H., E.S.-S., M.R., S.E.B., H.F.R., M.A.H., L.S., D.K., N.v.W., and J.K.; Antibody reactivity testing: H.-C.K.; Serological assays and neutralization testing: V.M.C., M.L.S., T.S., L.M.J., and C.D.; Protein production and crystallography: M.Y., W.Y., Y.H., and H.T.; Software: M.B., J.H., and S.M.R.; Resources: V.M.C., M.H., G.W., D.S., C.D., H.P., and I.A.W.; Writing – Original Draft: S.M.R., M.Y., H.-C.K., H.P., I.A.W., and J.K.; Writing – Review & Editing: all authors; Supervision: S.M.R., M.Y., H.-C.K., V.M.C., H.P., I.A.W., and J.K.

## Data and materials availability

X-ray coordinates and structure factors are deposited at the RCSB Protein Data Bank under accession codes 7S5P, 7S5Q and 7S5R. The amino acid sequences of the antibodies described in this study can be found in table S3. All requests for materials including antibodies, viruses, plasmids and proteins generated in this study should be directed to the corresponding authors. Materials will be made available for non-commercial usage.

## Materials and Methods

### Patient recruitment

All donors have given written informed consent and analyses were approved by the Institutional Review Board of Charité - Universitätsmedizin Berlin, corporate member of Freie Universität Berlin, Humboldt-Universität Berlin, and Berlin Institute of Health, Berlin (study protocol number EA1/258/18), the Institutional Review Board of the Faculty of Medicine at Ludwig-Maximilians-Universität (LMU) Munich, Germany (20-371), as well as the ethics committee of the Innsbruck Medical University (1167/2020). All donors in this study were tested positive for SARS-CoV-2 infection by quantitative PCR with reverse transcription (RT-qPCR) from nasopharyngeal swabs. Beta variant infection was confirmed using next-generation sequencing of swab material from the donor or the unequivocal contact person in the chain of infection. All donors were unvaccinated. Patient characteristics are described in table S1.

### Patient sample handling and single cell isolation

Peripheral blood mononuclear cells (PBMCs) were isolated by gradient centrifugation and then enriched for B cells by negative selection using a pan-B-cell isolation kit according to the manufacturer’s instructions (Miltenyi Biotec, 130-101-638).

His-tagged recombinant RBD Beta protein was produced in HEK cells (ACROBiosystems, SPD-C52Hp) and covalently labeled using CruzFluor488 (Santa Cruz Biotechnology, sc-362617) according to the manufacturer’s instructions. Separately, the same protein was labeled using its His-tag by incubating the antigen with an Alexa Fluor 647-conjugated anti-His-antibody (R&D Systems, IC0501R) for 30 minutes at room temperature at a 2:1 ratio (RBD molecules:IgG molecules). Ovalbumin (Sigma, A5503) was covalently labeled with PE-Cy7 (abcam, ab102903) according to the manufacturer’s instructions.

Using fluorescence-activated cell sorting (FACSAriaII SORP, BD Biosciences), we then sorted single RBD-Beta-double-positive 7AAD-Ovalbumin-CD19^+^CD27^+^ memory B cells (MBCs) into 96-well PCR plates. Staining was performed on ice for 25 minutes in PBS with 1 mM EDTA, 1:100 human IgG (1 mg/ml) as FcR block and 2 % FCS using the following staining reagents: 7-AAD 1:400 (Thermo Fisher Scientific), CD19-BV786 1:20 (clone SJ25C1, BD Biosciences, 563326), CD27-PE 1:5 (clone M-T271, BD Biosciences, 555441), Ovalbumin-PECy7 at 2 μg/ml, RBD-Beta-CruzFluor488 at 1 μg/ml (RBD concentration) and RBD-Beta/Anti-His-AF647 at 1 μg/ml (RBD concentration).

### Recombinant mAb generation

The mAbs were generated following our established protocols (*19*) with modifications as indicated. We used a nested PCR strategy to amplify the variable domains of the immunoglobulin heavy and light chain genes from single cell cDNA and analyzed their sequences with the aBASE module of Brain Antibody Sequence Evaluation (BASE) software (*32*). Pairs of functional Ig genes were PCR-amplified using specific primers with Q5 Polymerase (NEB). PCR-product and linearized vector containing the constant part of IgG1 heavy or kappa/lambda light chain sequences respectively were assembled using Gibson cloning with HiFi DNA Assembly Master Mix (NEB). Cloning was considered successful when sequence identity was >99.5% as verified by the cBASE module of BASE software. For mAb expression, human embryonic kidney cells (HEK293T) were transiently transfected with matching Ig heavy and light chains. The day after transfection, the supernatant was discarded and cells were supplemented with fresh medium. Six days later, mAb containing cell culture supernatant was harvested. Ig concentrations were determined and supernatants were used for reactivity and neutralization screening, if the Ig concentration was higher than 10 μg/ml. For biophysical characterization assays, supernatants were purified using Protein G Sepharose beads (GE Healthcare), dialyzed against PBS, and sterile-filtered using 0.2 μm filter units (GE Healthcare).

### mAb sequence analysis, clonotype analysis and data visualization

Sequence analysis, including gene usage, CDR3 length, and number of somatic hypermutations, was performed using our previously published script collection BASE (*32*). To identify antibodies which share the same V and J genes on both Ig heavy and light chains and thus are considered to be one clonotype, we used an in-house R script. After identification of public clonotypes, they were plotted in a Circos plot using the R package circlize (*33*). Our newly acquired dataset was compared to all previously published RBD mAbs included in the CoV-AbDab database, retrieved on 2021-06-16. As the CoV-AbDab includes SARS-CoV-2 mAbs from other sources than humans, and against other epitopes than the RBD, the following selection criteria were used (nomenclature like in CoV-AbDab): Binds to: SARS-CoV-2, Protein + Epitope: RBD, Origin: B-cells (human). 1157 mAbs fulfilled these criteria as human RBD mAbs, none of which were derived from studies of patients where infection with a VOC was reported. All 1037 mAbs for which information on V-J gene usage was available for both heavy and light chain were included in the clonotype analysis and VH gene usage analysis in Fig. 2.

### Diagnostic antibody testing

Initial serological testing of patient samples was performed using a solid phase immunoassay (SeraSpot®Anti-SARS-CoV-2 IgG, Seramun Diagnostica GmbH, Heidesee, Germany). Briefly, on the bottom of each well, SARS-CoV-2 recombinant antigens (nucleocapsid, non-VOC full spike, non-VOC S1 domain, non-VOC RBD) and controls are printed in an array format as spots. Serum samples were diluted 1:101 in sample dilution buffer, added to the wells, and incubated for 30 minutes at room temperature. After washing, bound antibodies were detected by incubation with horseradish peroxidase (HRP)-labeled anti-human IgG for 30 minutes at room temperature. After washing again, 3′3,5,5-tetramethylbenzidine (TMB) solution was added to each well and incubated in the dark for 30 minutes at room temperature. Subsequently, the solution was removed and color intensity of immune complexes formed at the site of each antigen spot was measured using a SpotSight®plate scanner. Color intensity correlates to the antibody concentration and results were calculated as signal-to-cutoff ratios using the internal cutoff control.

Second, the presence of SARS-CoV-2 S1-specific antibodies was analyzed using a commercially available anti-SARS-CoV-2-S1 IgG ELISA (EUROIMMUN Medizinische Labordiagnostika AG, Lübeck, Germany) according to the manufacturer’s instructions. Serum samples were diluted 1:101. The optical density (OD) at 450nm was measured and OD ratios were calculated by dividing this value by the OD of the kit-included calibrator.

Additionally, we applied a modified solid phase immunoassay (Seramun Diagnostica GmbH, Heidesee, Germany) as described above, which additionally contained SARS-CoV-2 RBD-VOCs (InVivo BioTech Services GmbH, Hennigsdorf, Germany).

### Surrogate virus neutralization test

Neutralizing capacity of patients’ sera was assessed by a surrogate virus neutralization test (cPass Assay, Medac, Wedel, Germany) according to the manufactureŕs instructions and as described previously (*34*). Briefly, serum samples, positive and negative controls were diluted 1:10 with sample dilution buffer, mixed 1:1 with non VOC HRP-RBD or Beta HRP-RBD (Medac, Wedel, Germany) solution and incubated at 37°C for 30 minutes. Afterwards, the mixture was added to the hACE2-coated plate and incubated at 37°C for 15 minutes. After a washing step, TMB solution was added, and the plate was incubated in the dark at room temperature for 15 minutes. Stop solution was then added and the optical density at 450 nm was measured using a Tecan Infinite 200 PRO plate reader. For calculation of the relative inhibition of ACE2/RBD binding, the following formula was applied: Inhibition score (%) = (1 − OD value sample/OD value negative control) × 100%.

### RBD ELISA

Binding of mAbs to SARS-CoV-2 spike RBD or variants thereof was detected by ELISA as previously described (*19*). Briefly, HEK293T cell-secreted RBD-Fc fusion proteins composed of the RBD-SD1 component of the SARS-CoV-2 spike S1 subunit (amino acids 319-591) and the constant region of rabbit IgG1 heavy chain (Fc) were immobilized onto 96-well plates via anti-rabbit IgG (Dianova, 711-005-152). Human mAbs were applied and detected using HRP-conjugated anti-human IgG (Dianova, 709-035-149) and the HRP substrate 1-step Ultra TMB (Thermo Fisher Scientific, Waltham, MA).

All mAbs were initially screened at 10 ng/ml for binding to RBD Beta and non-VOC RBD to identify strong binders and to distinguish Beta-selective from cross-reactive antibodies. RBD Beta-selective mAbs were tested at 100 ng/ml for binding to RBD Beta, non-VOC RBD as well as RBD variants containing all other combinations of amino-acid differences at positions 417 (N or K), 484 (K or E) and 501 (Y or N). Cross-reactive mAbs that neutralized SARS-CoV-2 Beta were tested at 100 ng/ml for binding to RBD-Fc proteins derived from the four current SARS-CoV-2 variants of concern (VOCs) Alpha (lineage B.1.1.7; RBD N501Y and SD1 A570D), Beta (B.1.351; RBD K417N, E484K and N501Y), Gamma (P.1; RBD K417T, E484K and N501Y) and Delta (B.1.617.2; RBD L452R and T478K) (*35, 36*); from SARS-CoV-2 non-VOC; and from SARS-CoV (*19*). For each mAb, the absorbance values at 450 nm based on binding to the RBD variants were normalized to its value on RBD Beta. Means of the relative values were determined from data of at least two independent experiments.

RBD Alpha and RBD Beta were generated based on gene synthesis of the S1 RBD-SD1 regions (Eurofins Genomics). Mutations for single amino-acid changes and for RBD Gamma and RBD Delta were introduced by overlap extension PCR. All constructs were checked by Sanger sequencing (LGC Genomics).

### Plaque reduction neutralization test

To assess the neutralizing activity of serum samples and SARS-CoV-2 mAbs, we performed plaque reduction neutralization tests (PRNT) as described (*14*) using virus of Beta isolate (GISAID accession no: EPI_ISL_862149), non-VOC Munich isolate 984 (*14*) and Delta isolate (GISAID accession no EPI_ISL_2500366). In brief, Vero E6 cells (1.6 x105 cells/well) were seeded in 24-well plates and incubated overnight. Serum or mAbs were diluted in OptiPro and mixed 1:1 with 200 μL of the respective virus isolate solution containing 100 plaque forming units. The serum- or mAb-virus solutions were incubated on the Vero E6 cells for 1 hour at 37°C, then discarded, and cells were washed once with PBS and supplemented with 1.2% Avicel solution in DMEM. After three days of incubation, the supernatants were removed, the cells were fixed and inactivated using a 6% formaldehyde in PBS solution and then stained with crystal violet to count plaques. mAbs were diluted to 200 and 2000 ng/ml in mAb-virus solution for screening assays and in further dilutions for the determination of the IC50. Serum samples were serially diluted starting at 1:20. All dilutions were tested in duplicate.

### Surface Plasmon resonance

Anti-human IgG (Fc) capture antibody was covalenty immobilized on a C1 sensor chip. Purified mAbs were reversibly immobilized via the anti-human IgG capture surface. The RBD Beta protein (Acro Biosystems, SPD-C52Hp) was injected at different concentrations in a buffer consisting of 10 mM HEPES pH 7.4, 150 mM NaCl, 3 mM EDTA, 0.05% Tween 20, 0.1 mg/ml BSA. Ka, Kd and KD-values were determined using a monovalent analyte model. Recordings were performed on a Biacore T200 instrument at 25°C.

### Crystallization and structural determination

The RBD (residues 333-529) of the SARS-CoV-2 spike (S) protein (GenBank: QHD43416.1) and the Beta variant that carries three mutations on the RBD (K417N, E484K, and N501Y) were cloned into a customized pFastBac vector (*37*), and fused with an N-terminal gp67 signal peptide and C-terminal His6 tag. A recombinant bacmid DNA was generated using the Bac-to-Bac system (Life Technologies). Baculovirus was generated by transfecting purified bacmid DNA into Sf9 cells using FuGENE HD (Promega), and subsequently used to infect suspension cultures of High Five cells (Life Technologies) at an MOI of 5 to 10. Infected High Five cells were incubated at 28 °C with shaking at 110 r.p.m. for 72 h for protein expression. The supernatant was then concentrated using a 10 kDa MW cutoff Centramate cassette (Pall Corporation). The RBD protein was purified by Ni-NTA, followed by size exclusion chromatography, and buffer exchanged into 20 mM Tris-HCl pH 7.4 and 150 mM NaCl.

For expression and purification of the Fabs, heavy and light chains were cloned into phCMV3. The plasmids were transiently co-transfected into ExpiCHO cells at a ratio of 2:1 (HC:LC) using ExpiFectamine™ CHO Reagent (Thermo Fisher Scientific) according to the manufacturer’s instructions. The supernatant was collected at 14 days post-transfection. The Fabs were purified with a CaptureSelect™ CH1-XL Affinity Matrix (Thermo Fisher Scientific) followed by size exclusion chromatography.

CS23/B.1.351 RBD, CS44/COVA1-16/B.1.351 RBD, and CV07-287/COVA1-16/wild-type RBD complexes were formed by mixing each of the protein components at an equimolar ratio and incubated overnight at 4°C. COVA1-16 Fabs were used to assist with the crystal packing (*38*). Each complex was screened for crystallization using the 384 conditions of the JCSG Core Suite (Qiagen) and ProPlex screen (Molecular Dimensions) on either our robotic CrystalMation system (Rigaku) or an Oryx8 (Douglas Instruments) at Scripps Research. Crystallization trials were set-up by the vapor diffusion method in sitting drops containing 0.1 μl of protein and 0.1 μl of reservoir solution. Diffraction-quality crystals were obtained in the following conditions:

CS23/B.1.351 RBD (12.5 mg/ml): 1.6 M ammonium sulfate and 0.1 M bicine pH 9.0 at 20°C CS44/COVA1-16/B.1.351 RBD (12.0 mg/ml): 1.6 M ammonium sulfate and 0.1 M citric acid pH 4.0 at 20°C

CV07-287/COVA1-16/wild-type RBD (12.0 mg/ml): 1.6 M ammonium sulfate and 0.1 M bicine pH 9.0 at 20°C

All crystals appeared on day 3 and were harvested on day 7. Before flash cooling in liquid nitrogen for X-ray diffraction studies, crystals were equilibrated in reservoir solution supplemented the following cryoprotectants:

CS23/B.1.351 RBD: 20% ethylene glycol

CS44/COVA1-16/B.1.351 RBD: 20% glycerol

CV07-287/COVA1-16/wild-type RBD: 20% ethylene glycol

Diffraction data were collected at cryogenic temperature (100 K) at the Advanced Light Source on the beamline 5.0.1 and the Stanford Synchrotron Radiation Lightsource (SSRL) on Scripps/Stanford beamline 12-1, and processed with HKL2000 (*39*) (table S4). Structures were solved by molecular replacement using PHASER (*40*) with PDB 6W41. Iterative model building and refinement were carried out in COOT (*41*) and PHENIX (*42*), respectively (table S4). Epitope and paratope residues, as well as their interactions, were identified by accessing PISA at the European Bioinformatics Institute (http://www.ebi.ac.uk/pdbe/prot_int/pistart.html) (*43*) (table S5).

### Biolayer interferometry (BLI) competition assay

Competition assays were performed by biolayer interferometry (BLI) using an Octet Red instrument (FortéBio). IgGs were diluted with kinetic buffer (1x PBS, pH 7.4, 0.01% BSA and 0.002% Tween 20). After His-tagged RBD Beta was immobilized on anti-Penta His BLI sensors, sensors were first dipped into CS82 IgG (50 μg/ml), and then dipped into indicated IgG antibodies (12.5 μg/ml). Three replicates were performed for each BLI experiment.

### Statistical analyses

Area under the curve (AUC) calculations in Fig. 1 and all statistical analyses were performed using GraphPad Prism (9.2.0).

---

**Fig. S1.**
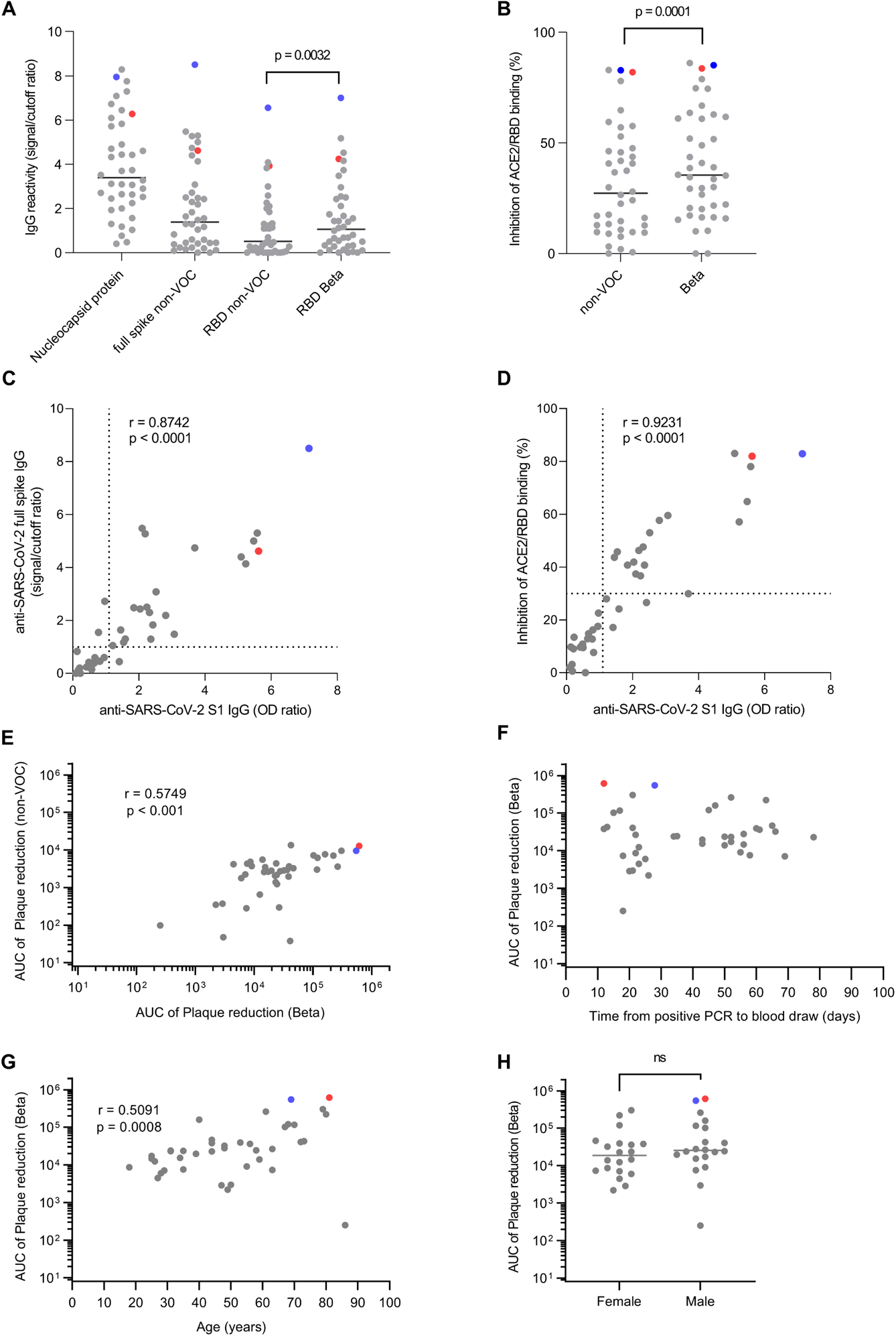
Polyclonal antibodies from individuals after infection with SARS-CoV-2 Beta. **(A)** IgG reactivity against indicated antigens was measured by SeraSpot Anti-SARS-CoV-2 IgG assay. Statistical analysis was performed using a Friedman test and Dunn’s multiple comparison test. Only the p-value for comparison of non-VOC RBD vs. RBD Beta is shown. Horizontal bars indicate median values. **(B)** ACE2 interaction with indicated RBD was determined using the cPass Surrogate Neutralization Assay. Statistical analysis was performed using a Wilcoxon matched-pairs signed rank test. Horizontal bars indicate median values. Values below zero were set to zero, indicating no inhibition. **(C-D)** Anti-SARS-CoV-2 S1 IgG measured by Euroimmun S1-ELISA is plotted against **(C)** anti-SARS-CoV-2 full non-VOC spike IgG measured by SeraSpot IgG assay and against **(D)** inhibition of ACE2/non-VOC RBD binding measured by cPass Surrogate Neutralization assay. Statistical analysis was performed by two-tailed Spearman’s correlation. Dashed lines indicate cutoffs defined by the assays’ manufacturers, which have been validated for non-VOC sera. **(E)** Correlation of neutralization activity against SARS-CoV-2 Beta and SARS-CoV-2 wildtype measured as AUC of PRNT is shown. Statistical analysis was performed by two-tailed Spearman’s correlations. **(F)** Time between first positive PCR test and sample collection in days is plotted against the AUC of PRNT against SARS-CoV-2 Beta. Statistical analysis performed by two-tailed Spearman’s correlations showed no statistically significant correlation. **(G)** Patient age is plotted against AUC of PRNT against SARS-CoV-2 Beta. Statistical analysis was performed by two-tailed Spearman’s correlations. **(H)** AUC of PRNT is plotted against SARS-CoV-2 Beta for men (n = 20) and women (n = 20) Statistical significance was determined using two-tailed Mann–Whitney U-tests. **(E-H)** AUC calculation is based on authentic virus PRNT curves (shown in Fig. 1, A and B). Patients SA1 and SA2 are highlighted in red and blue, respectively.

**Fig. S2.**
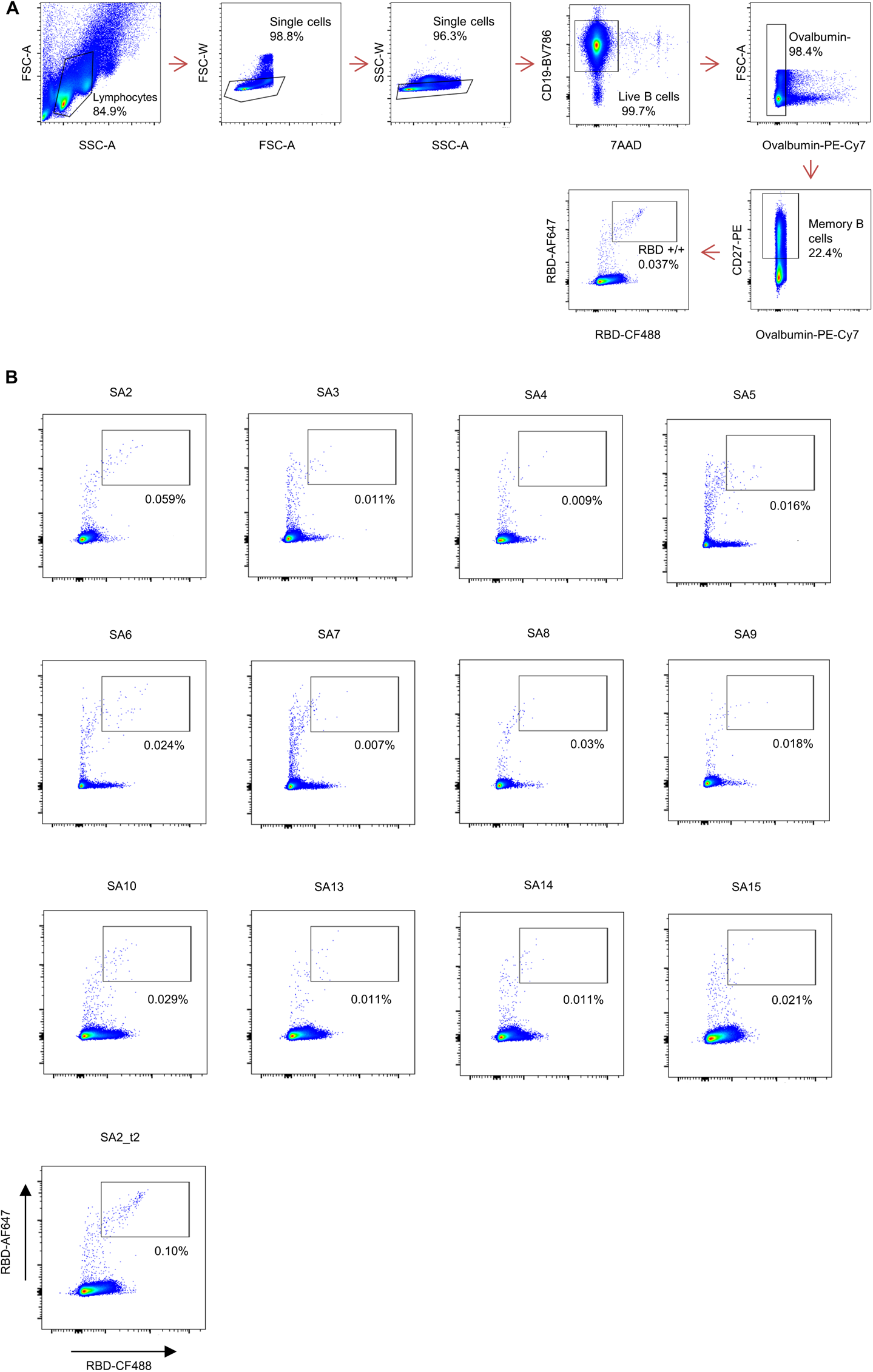
Gating strategy used for single cell sorting of RBD-reactive memory B cells. **(A)** Gating was on singlets that were CD19^+^7AAD^-^Ovalbumin^-^CD27^+^. Sorted cells were RBD-AF647^+^/RBD-CF488^+^. **(B)** Flow cytometry showing the percentage of RBD-double-positive memory B cells of indicated patients. Patient SA2 donated blood twice, and the analysis from the second time point is denoted by SA2_t2.

**Fig. S3.**
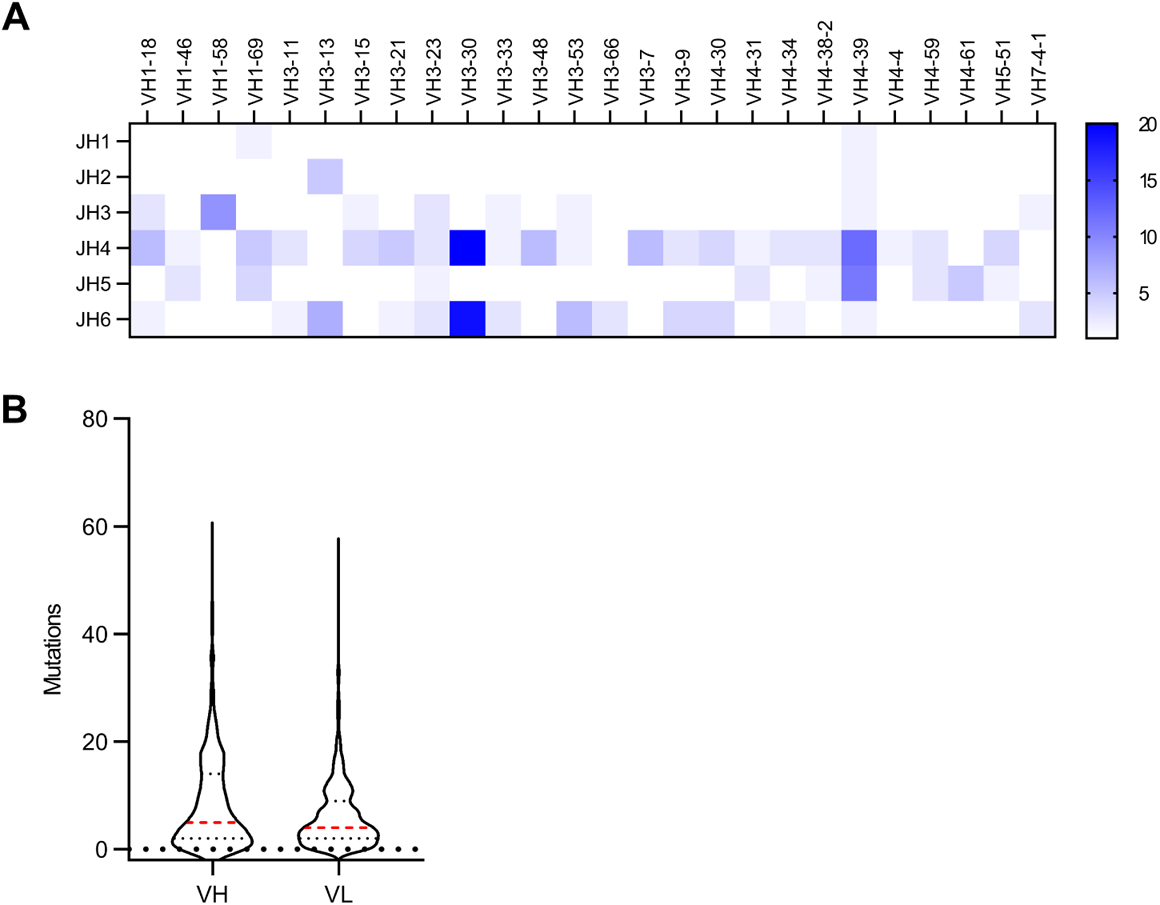
Sequence characteristics of Beta-elicited monoclonal antibodies in this study. **(A)** Pairing of indicated VH genes with JH gene families of 289 IgG mAbs are shown in absolute numbers. Only VH genes with 4 or more occurrences are shown. **(B)** Violin plots of somatic nucleotide mutations in the IGVH and IGVL genes in antibodies obtained from all donors. Red bars indicate mean.

**Fig. S4.**
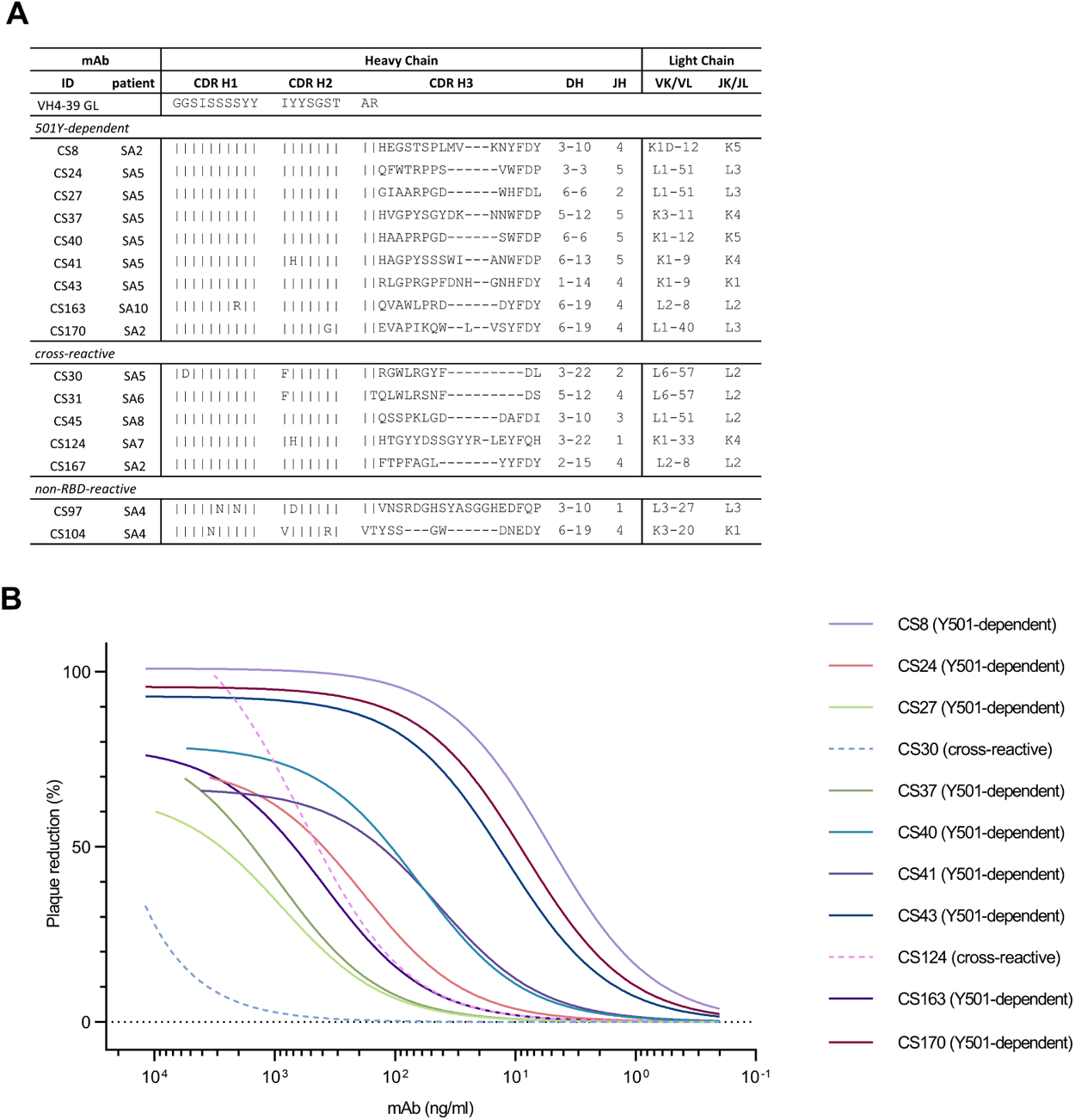
Neutralization of Beta-specific VH4-39 antibodies from multiple patients infected with SARS-CoV-2 Beta. **(A)** Comparison of sequence features of all expressed VH4-39 mAbs in this study. All antibodies which were expressed are shown in this panel. **(B)** Fitted curves of VH4-39 mAbs show dose-dependent neutralization against authentic SARS-CoV-2 Beta variant virus.

**Fig. S5.**
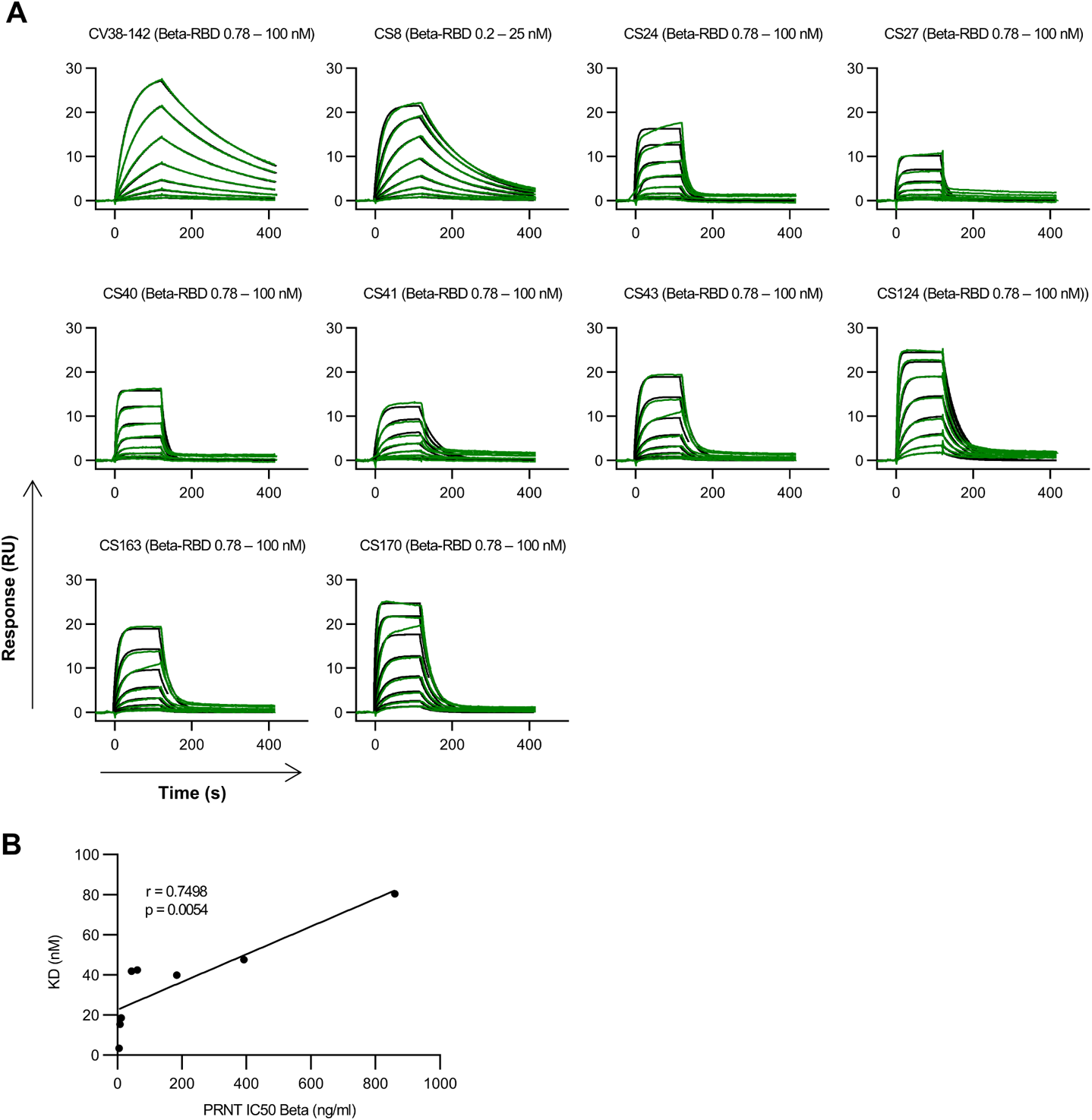
Binding kinetic measurements of Beta-RBD to VH4-39 mAbs. **(A)** Binding kinetics of Beta-RBD to indicated mAbs were modeled (black) from multi-cycle surface plasmon resonance (SPR) measurements (green). The fitted monovalent analyte model is shown. For CS24, CS27 and CS41, there was a second phase after the fast dissociation impeding the quality of the monovalent analyte model. All measurements are performed using serial 2-fold dilutions of SARS-CoV-2 Beta-RBD-His on immobilized mAbs. **(B)** Correlation of neutralization and affinity of monoclonal VH4-39 Y501-specific antibodies. IC_50_ was determined from neutralization curves shown in fig. S4B. Affinity of monoclonal antibodies to Beta RBD was determined by the fitted analyte model shown in (**A**). Statistical analysis was performed by a simple linear regression.

**Fig. S6.**
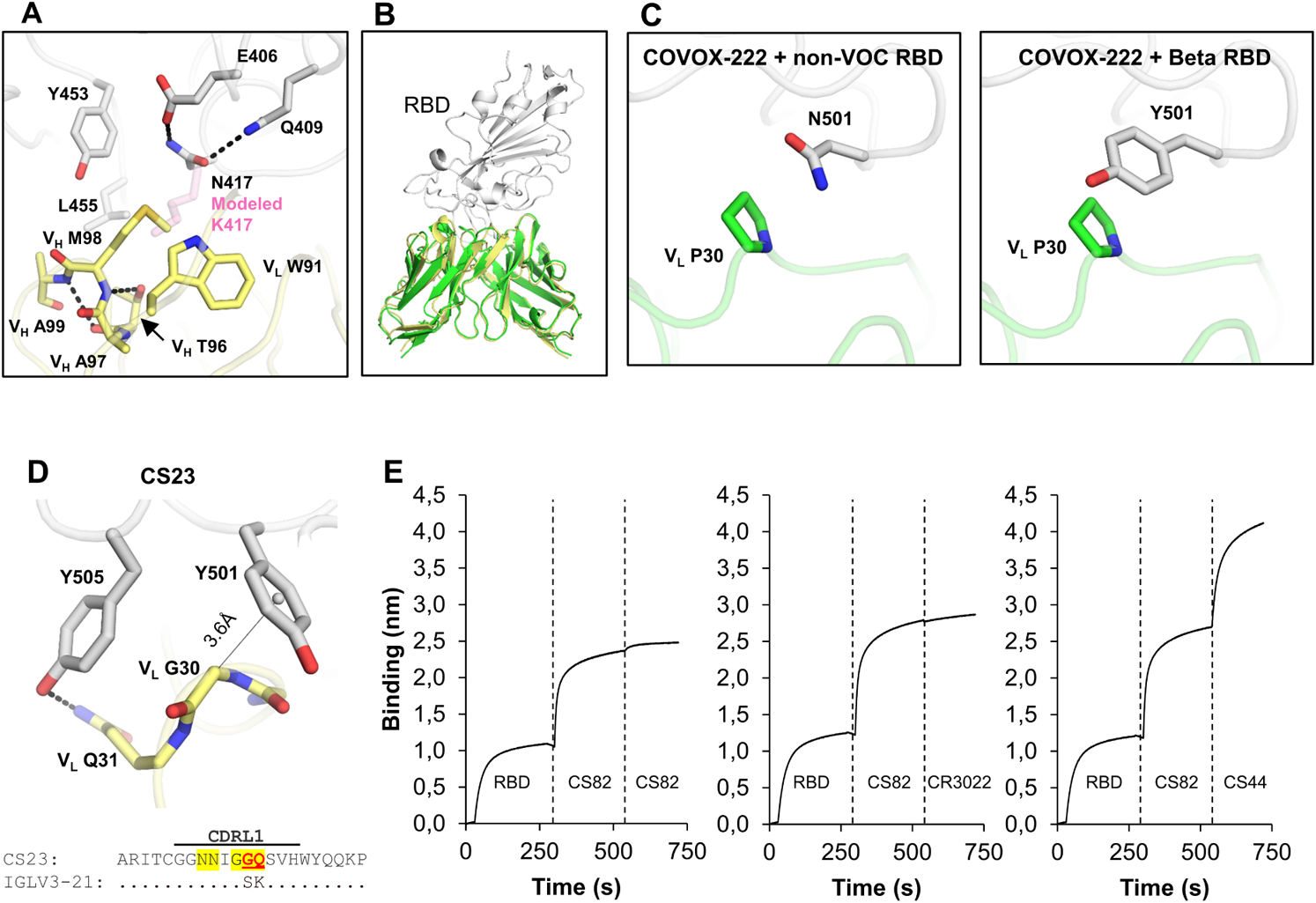
Structural and binding analysis of VH3-53 mAbs that bind to Beta RBD. The RBD is shown in white. Hydrogen bonds are represented by black dashed lines. **(A)** Structure of the CDR H3 of CS23 (yellow). A modeled side chain of K417 is shown as transparent pink sticks, which would be unfavorable for binding to CS23, where VH M98 occupies this pocket. **(B)** COVOX-222 and CS23 adopt the same binding mode. The crystal structure of COVOX-222 (green) in complex with RBD (Beta) was superimposed onto the structure of CS23 (yellow) in complex with RBD (Beta). Only the variable domains of the antibodies are shown for clarity. **(C)** Structures of COVOX-222 (green) in complex with non-VOC RBD (left, PDB 7NX6) and Beta RBD (right, PDB 7NXA). VL P30 and Y501 in the Beta RBD forms a π-π interaction. **(D)** CDR L1 of CS23 (yellow) interacts with the Beta RBD (white). Hydrogen bonds are represented by black dashed lines. The distance between V_L_ G30-Cα and the benzene ring center of RBD-Y501 (white sphere) is represented by a thin black solid line. Sequence alignment between the light chain of CS23 and its germline sequence is shown at the bottom, where identical residues are represented by dots. Paratope residues (defined as buried surface area > 0 Å^2^) are highlighted in yellow, and somatically mutated residues shown in red letters. **(E)** Biolayer interferometry competition assay between antibodies. The biosensor was first loaded with SARS-CoV-2 RBD, followed by two binding events: 1) CS82 IgG; 2) CS82, CR3022, or CS44 IgGs. Representative results of three replicates for each experiment are shown.

**Table S1.** Clinical characteristics and neutralization of serum by patients included in this study.

| Patient | Sex | Age | Date of first positive PCR test | Days from positive PCR to blood draw | Symptoms | Hospitalization | AUC PRNT Beta | AUC PRNT non-VOC (Munich isolate) | ACE2/RBD inhibition non-VOC (percent)<sup>a</sup> | ACE2/RBD inhibition Beta (percent)<sup>a</sup> |
| --- | --- | --- | --- | --- | --- | --- | --- | --- | --- | --- |
| SA1 | m | 81 | 22.01.2021 | 12 | yes | yes | 614037 | 12807 | 82.0 | 83.7 |
| SA2 | m | 69 | 27.01.2021 | 28 | yes | yes | 547486 | 9557 | 82.9 | 85.2 |
| SA3 | f | 49 | 29.01.2021 | 26 | yes | no | 2210 | 351,1 | 7.65 | 16.4 |
| SA4 | f | 18 | 02.02.2021 | 22 | yes | no | 8601 | 4879 | 47.6 | 40.9 |
| SA5 | f | 27 | 03.02.2021 | 23 | yes | no | 4466 | 4173 | 53.0 | 35.3 |
| SA6 | f | 28 | 01.02.2021 | 25 | yes | no | 6061 | 1794 | 40.7 | 33.7 |
| SA7 | f | 26 | 03.02.2021 | 23 | yes | no | 12492 | 654,1 | 16.2 | 30.0 |
| SA8 | m | 63 | 04.02.2021 | 22 | yes | no | 26705 | 296,6 | 9.7 | 10.3 |
| SA9 | f | 63 | 08.02.2021 | 18 | yes | no | 7399 | 281,2 | 9.0 | 16.0 |
| SA10 | m | 67 | 22.02.2021 | 15 | yes | no | 101879 | 7195 | 78.1 | 74.8 |
| SA11 | m | 72 | 16.02.2021 | 21 | no | no | 41076 | 38 | 9.5 | 22.2 |
| SA12 | m | 86 | 19.02.2021 | 18 | yes | no | 250 | 98,8 | 0.6 | 10.1 |
| SA13 | f | 79 | 16.02.2021 | 21 | yes | no | 304474 | 9535 | 59.5 | 62.9 |
| SA14 | m | 50 | 16.02.2021 | 21 | yes | no | 2982 | 47,5 | 13.4 | 17.2 |
| SA15 | f | 47 | 17.02.2021 | 20 | yes | no | 2879 | 372,7 | 3.1 | 0 |
| SA17 | f | 68 | 18.01.2021 | 45 | yes | yes | 119883 | 6225 | 46.3 | 74.5 |
| SA19 | m | 70 | 16.02.2021 | 17 | yes | yes | 117433 | 3033 | 57.1 | 61.0 |
| SA20 | f | 44 | 24.02.2021 | 12 | yes | yes | 38354 | 1959 | 29.9 | 26.7 |
| SA21 | f | 59 | 19.01.2021 | 50 | no | no | 13887 | 5602 | 37.4 | 60.9 |
| SA23 | f | 35 | 19.01.2021 | 50 | yes | no | 23737 | 4387 | 17.2 | 35.7 |
| SA24 | f | 44 | 04.01.2021 | 65 | yes | no | 46577 | 3262 | 57.7 | 63.7 |
| SA25 | m | 39 | 26.01.2021 | 43 | yes | no | 19655 | 2838 | 12.9 | 15.3 |
| SA26 | m | 58 | 03.02.2021 | 35 | yes | no | 24506 | 1246 | 40.7 | 49.0 |
| SA27 | m | 34 | 26.01.2021 | 43 | yes | no | 15453 | 3490 | 22.8 | 20.6 |
| SA28 | m | 61 | 17.01.2021 | 52 | yes | yes | 263808 | 3634 | 64.8 | 78.9 |
| SA29 | m | 55 | 14.01.2021 | 55 | yes | no | 9084 | 3722 | 2.0 | 0 |
| SA30 | f | 29 | 31.12.2020 | 69 | yes | no | 7079 | 2242 | 0 | 20.2 |
| SA31 | m | 40 | 23.01.2021 | 47 | no | no | 160344 | 7747 | 83.0 | 86.1 |
| SA32 | f | 80 | 07.01.2021 | 63 | yes | yes | 223456 | 7066 | 45.8 | 52.0 |
| SA33 | f | 25 | 14.01.2021 | 56 | no | no | 14762 | 2588 | 10.8 | 16.4 |
| SA34 | f | 56 | 09.01.2021 | 61 | yes | no | 36553 | 2994 | 41.9 | 39.1 |
| SA35 | m | 31 | 18.01.2021 | 52 | yes | no | 23301 | 1388 | 27.9 | 38.7 |
| SA36 | m | 25 | 18.01.2021 | 52 | yes | no | 17300 | 2663 | 43.7 | 34.5 |
| SA37 | f | 53 | 11.01.2021 | 60 | yes | no | 39374 | 3673 | 24.2 | 51.7 |
| SA38 | f | 44 | 27.12.2020 | 78 | yes | no | 22795 | 2083 | 17.6 | 29.6 |
| SA39 | m | 31 | 09.02.2021 | 34 | yes | no | 24121 | 2182 | 12.8 | 21.8 |
| SA40 | m | 48 | 18.01.2021 | 56 | yes | no | 27711 | 2745 | 36.7 | 66.9 |
| SA41 | f | 48 | 08.01.2021 | 66 | yes | no | 32266 | 2855 | 26.6 | 61.8 |
| SA42 | m | 35 | 16.01.2021 | 58 | yes | no | 7638 | 4355 | 14.8 | 29.4 |
| SA43 | m | 73 | 03.03.2021 | 13 | yes | yes | 42477 | 13482 | 9.8 | 43.6 |

<sup>a</sup> Values below zero were set to zero, indicating no inhibition.

**Table S2.**
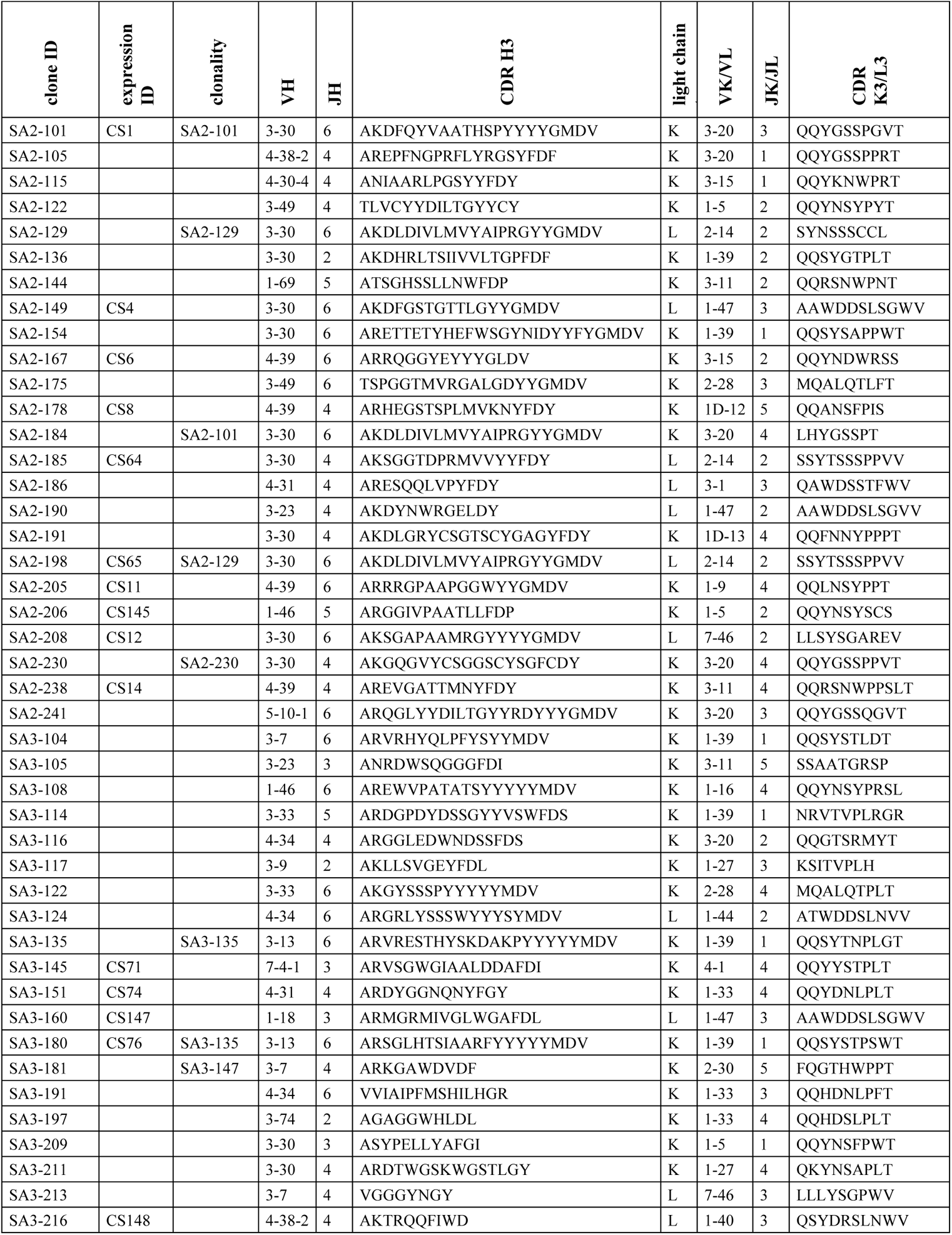
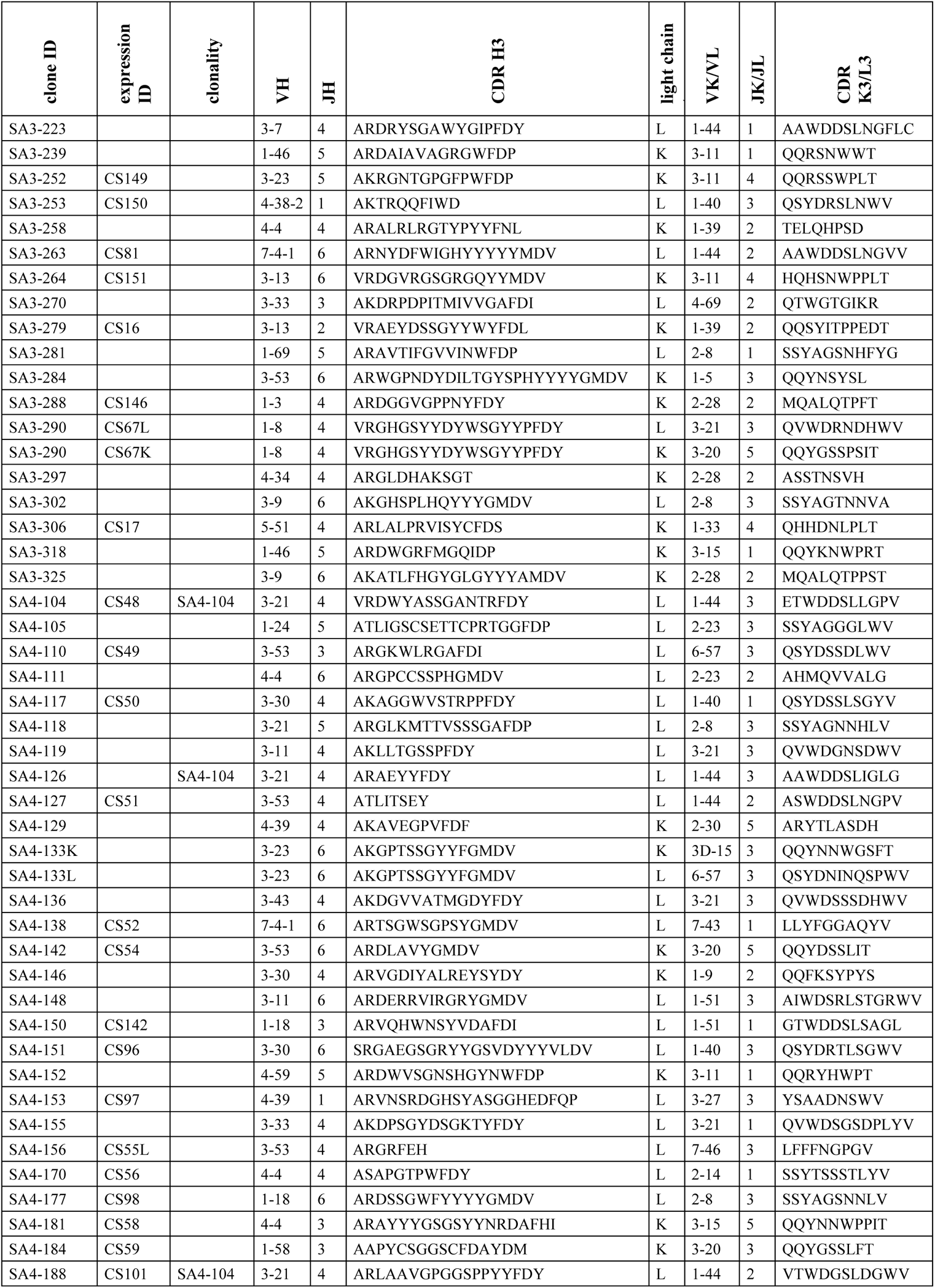
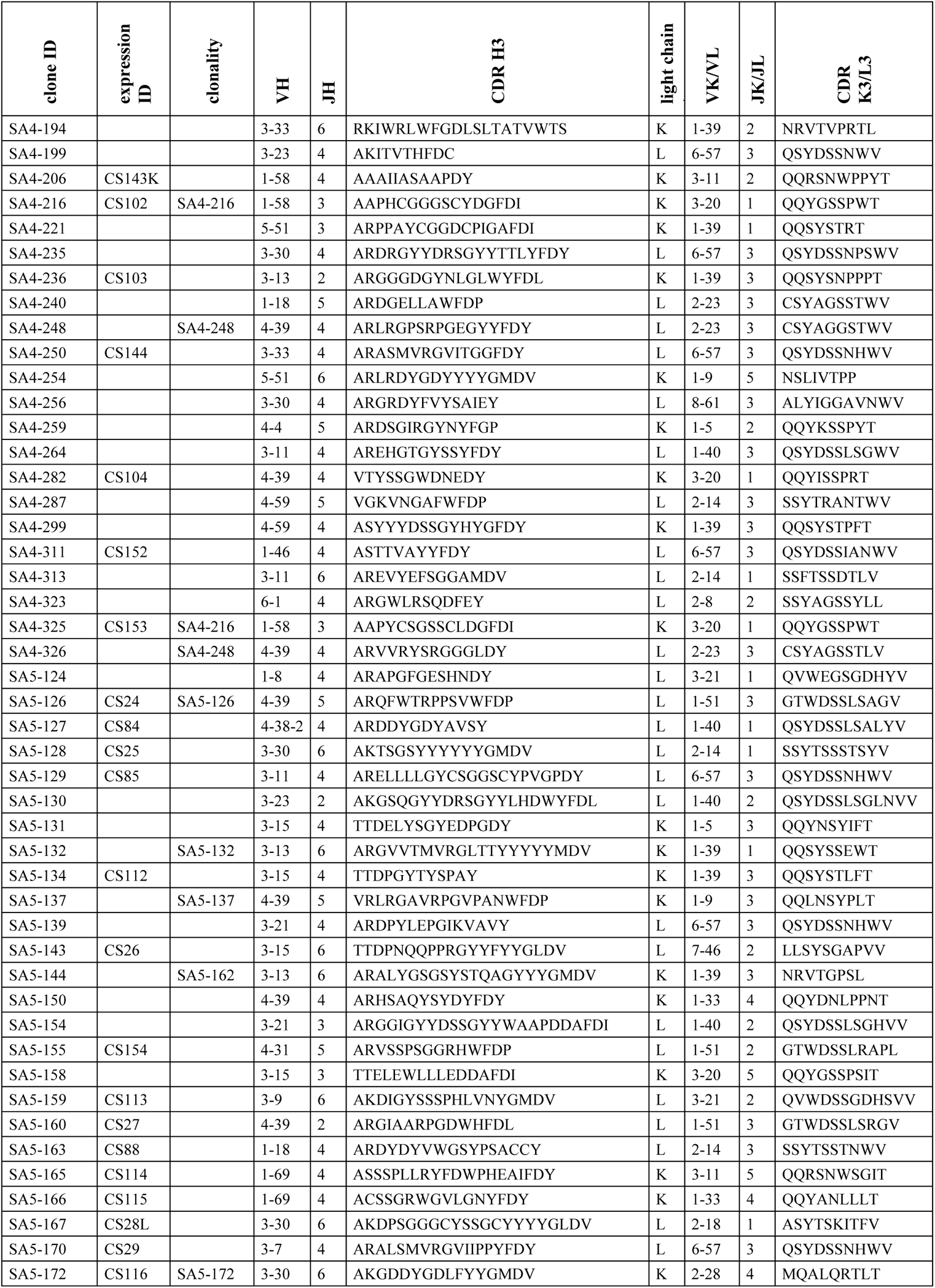
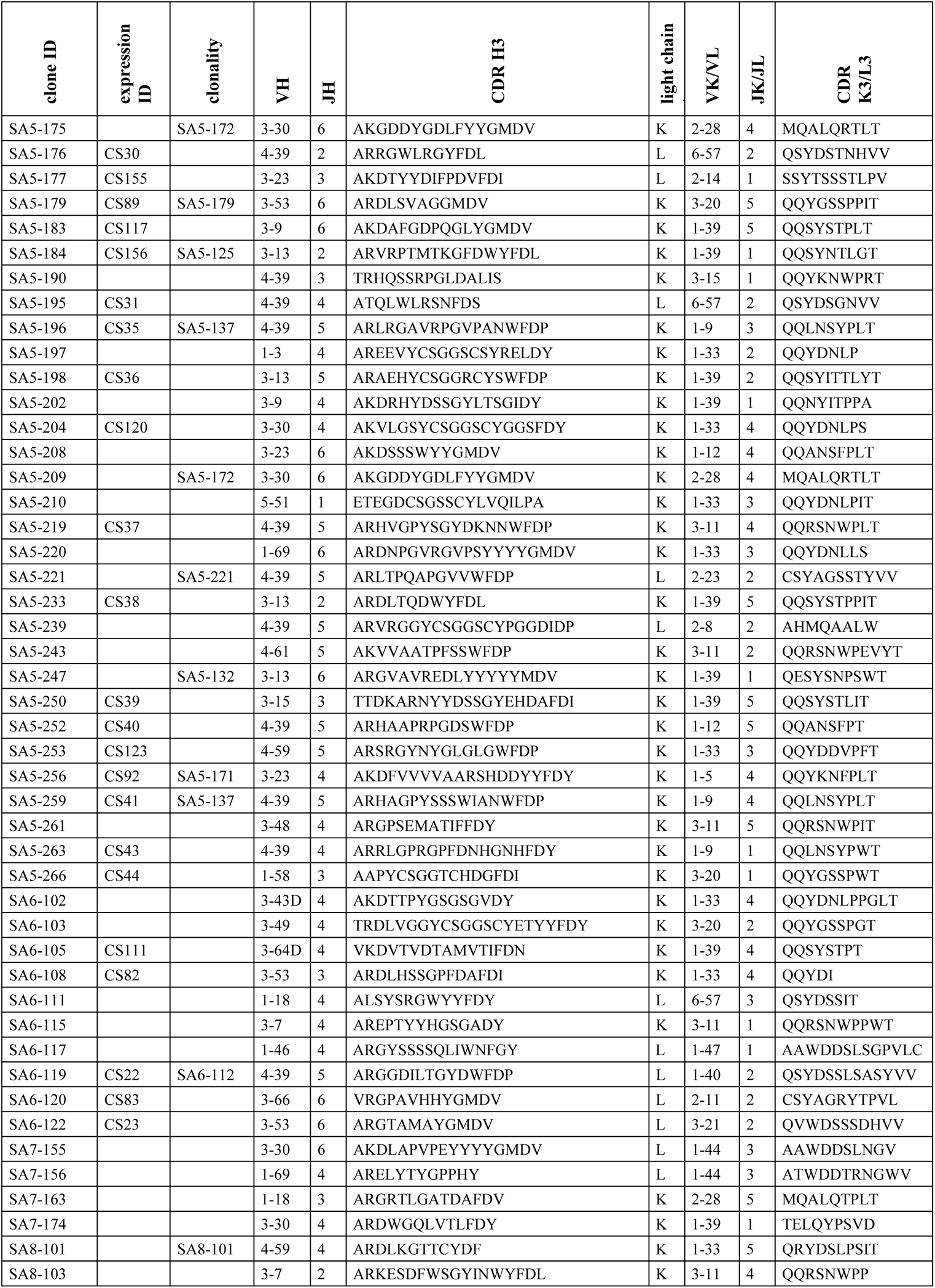
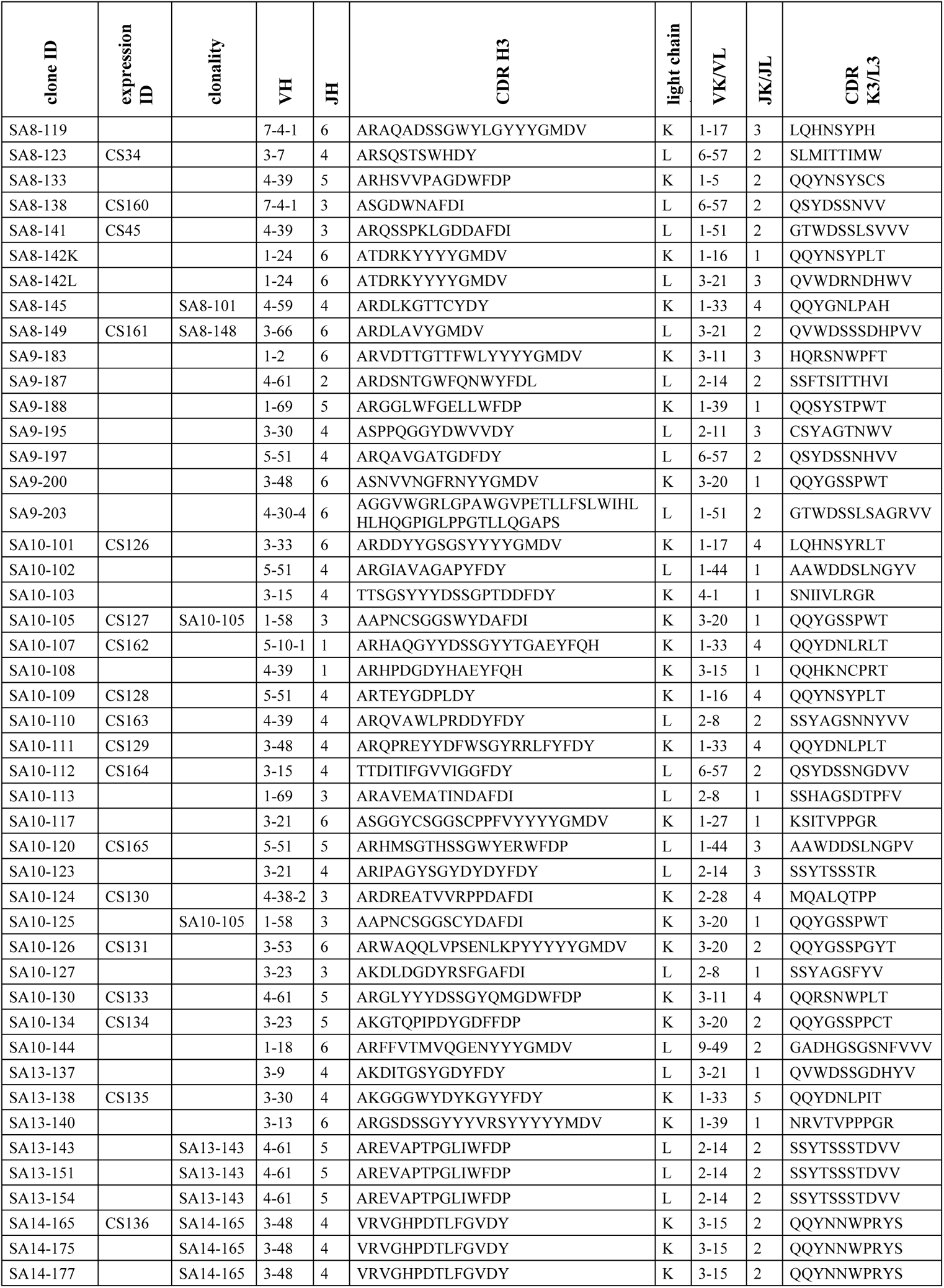
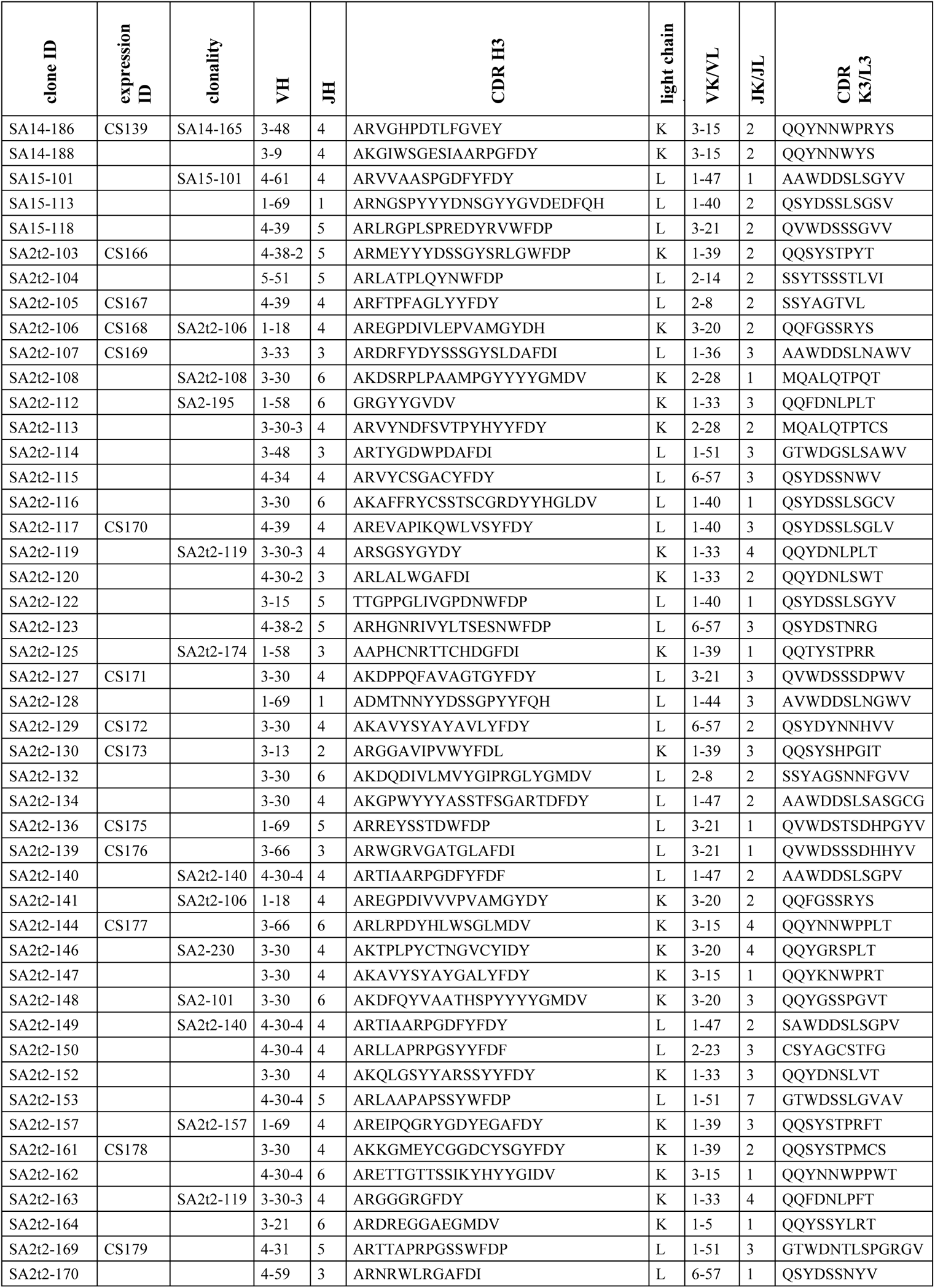
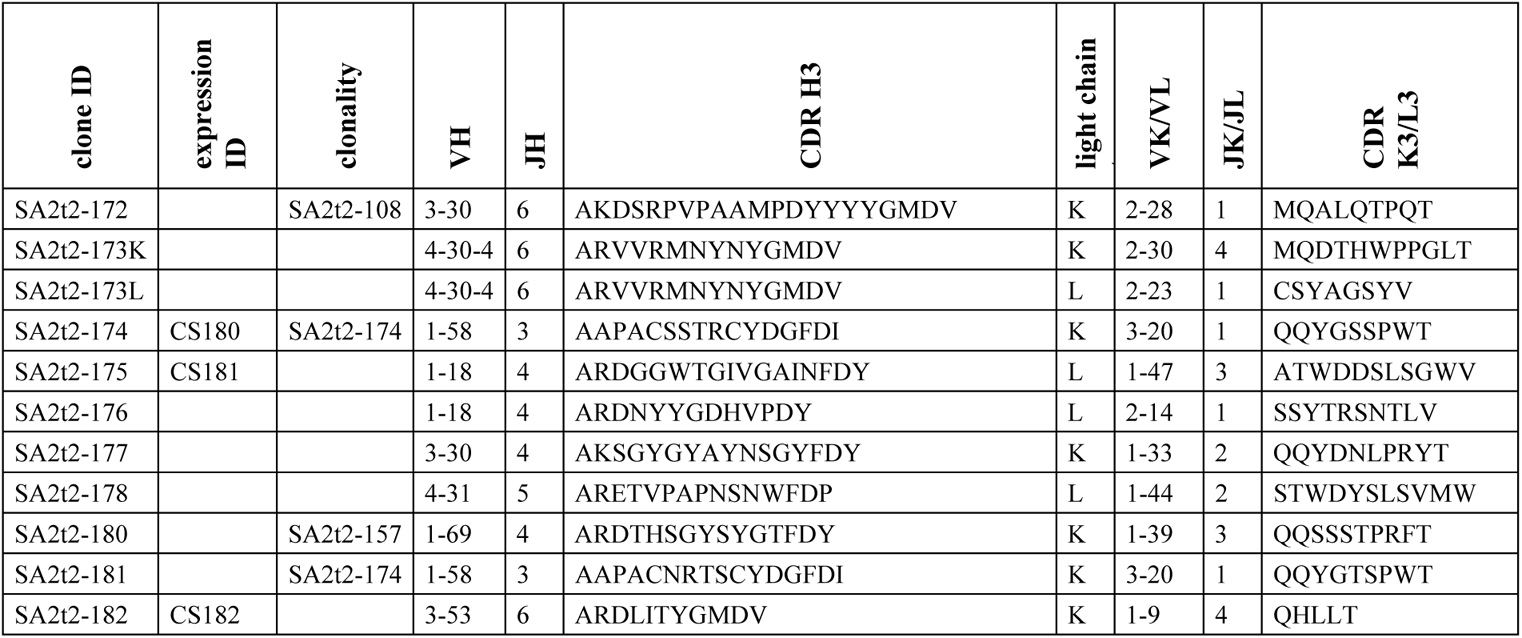
Sequence features of monoclonal antibodies in this study.

**Table S3.**
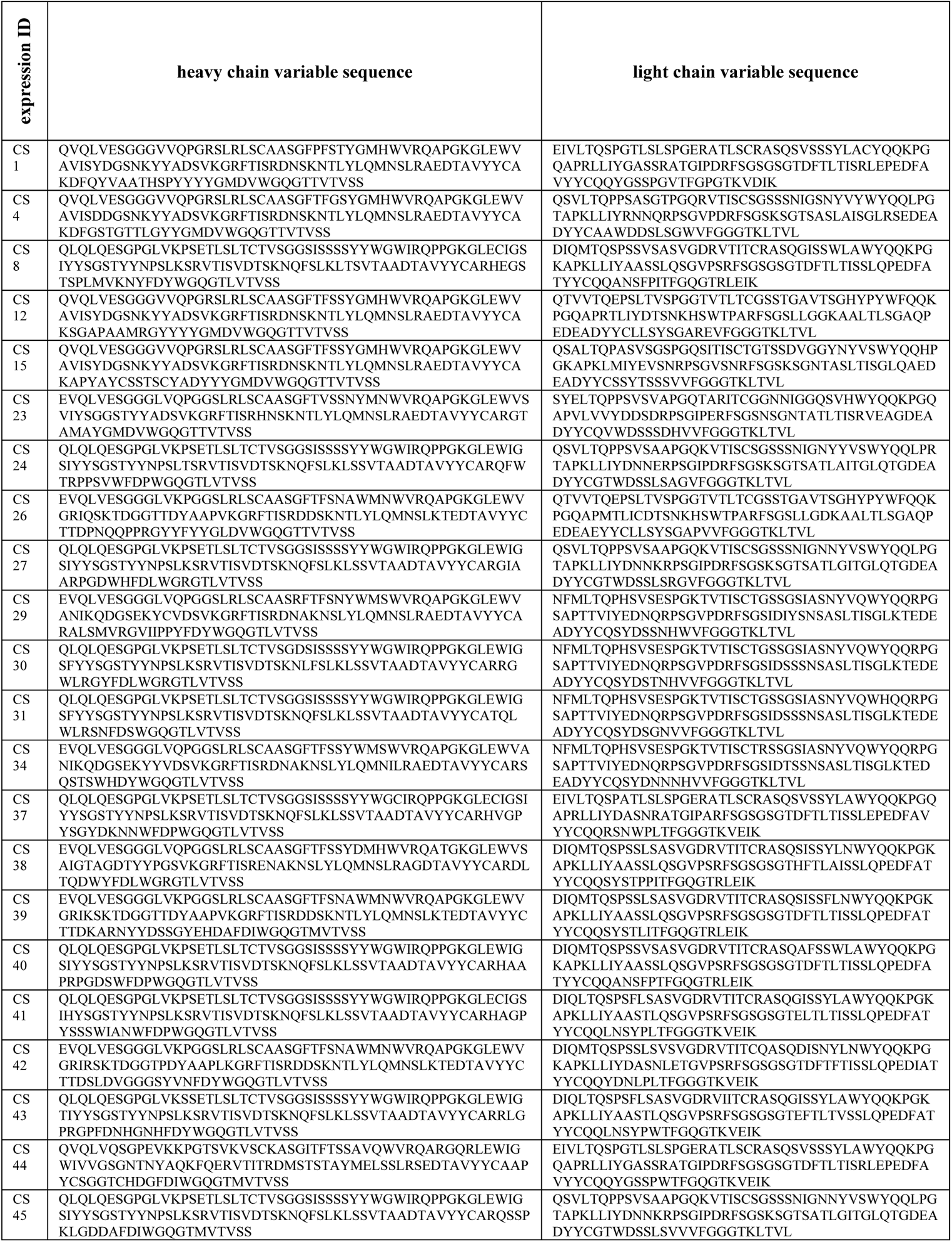
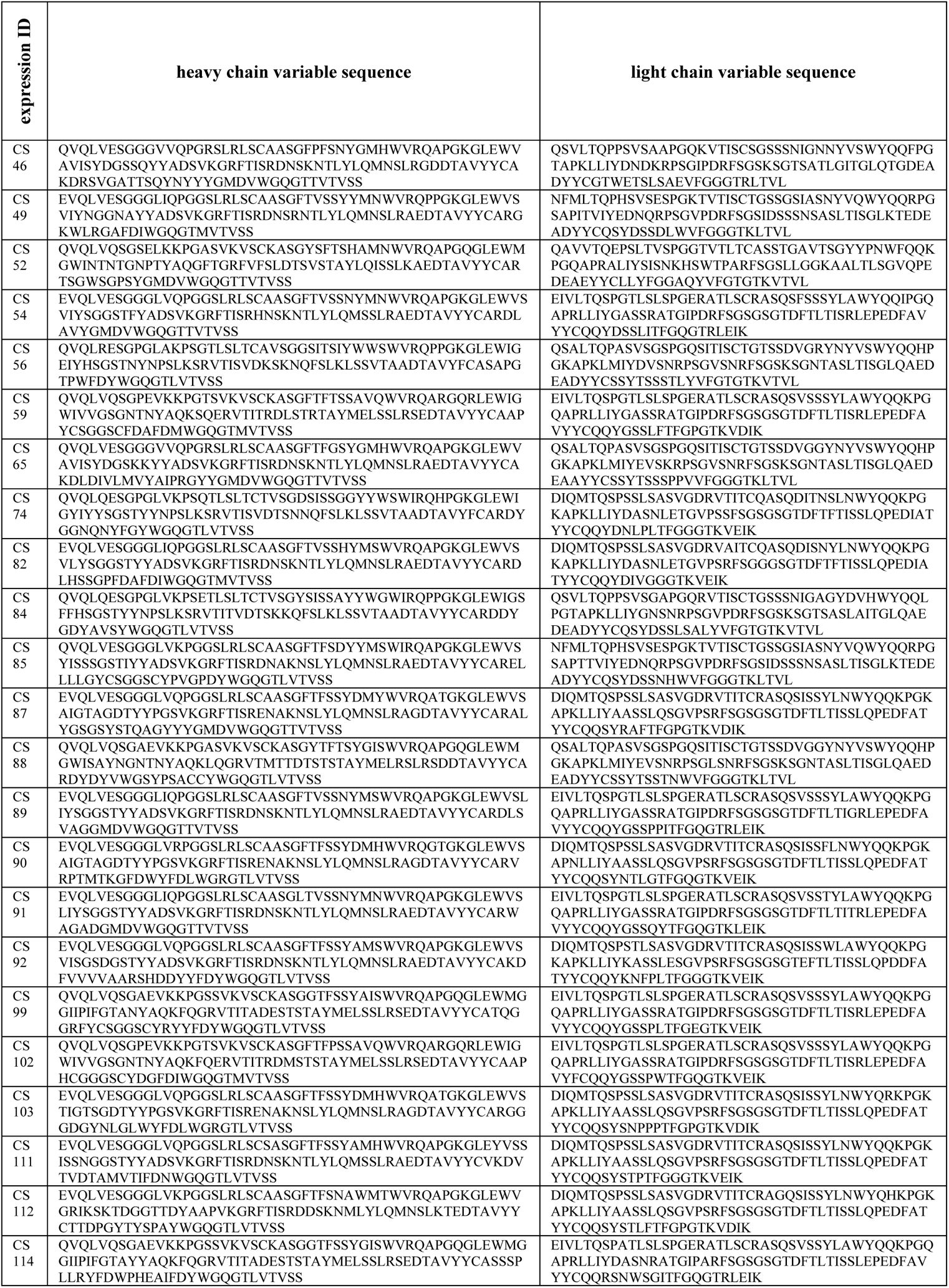
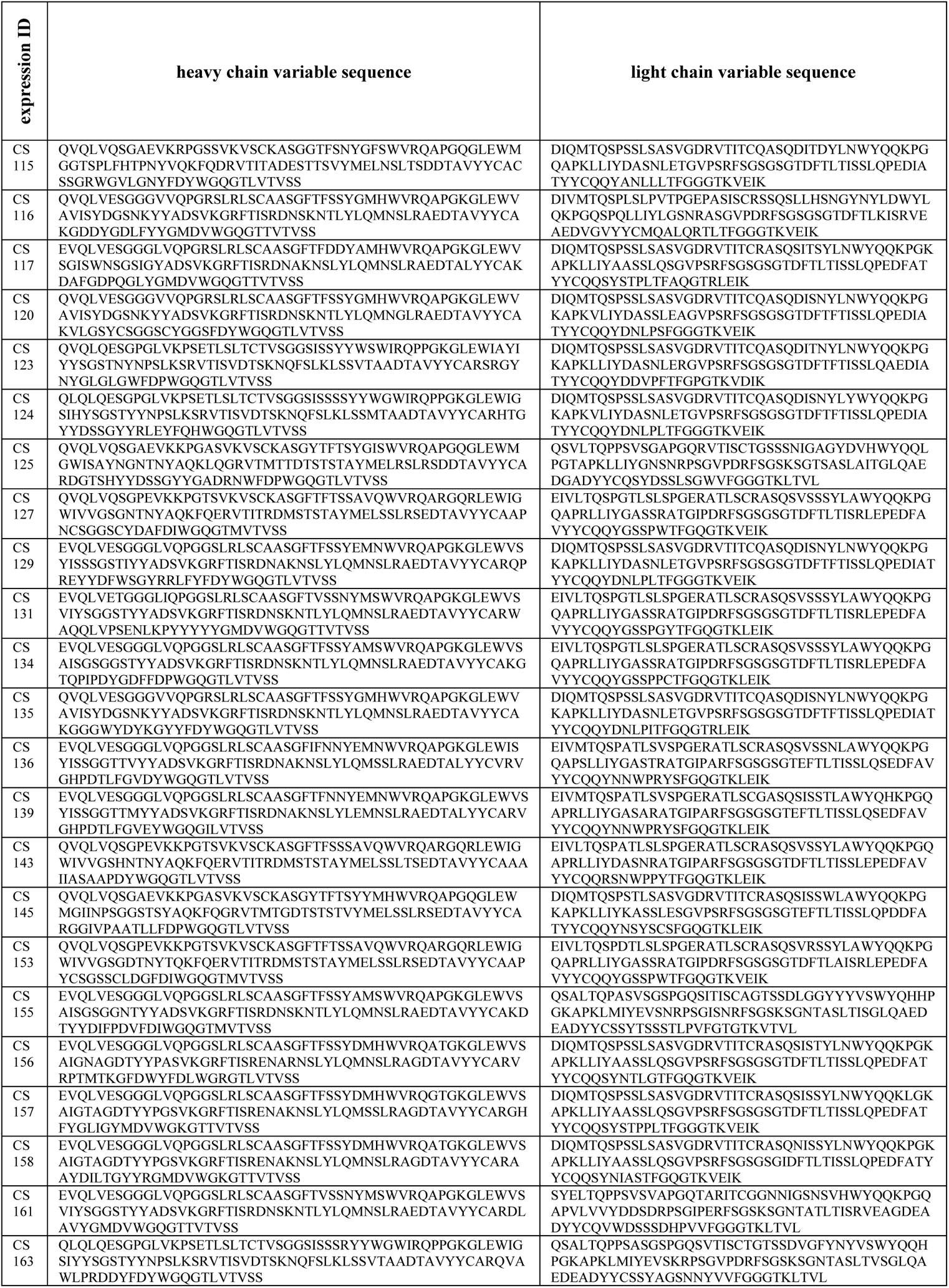
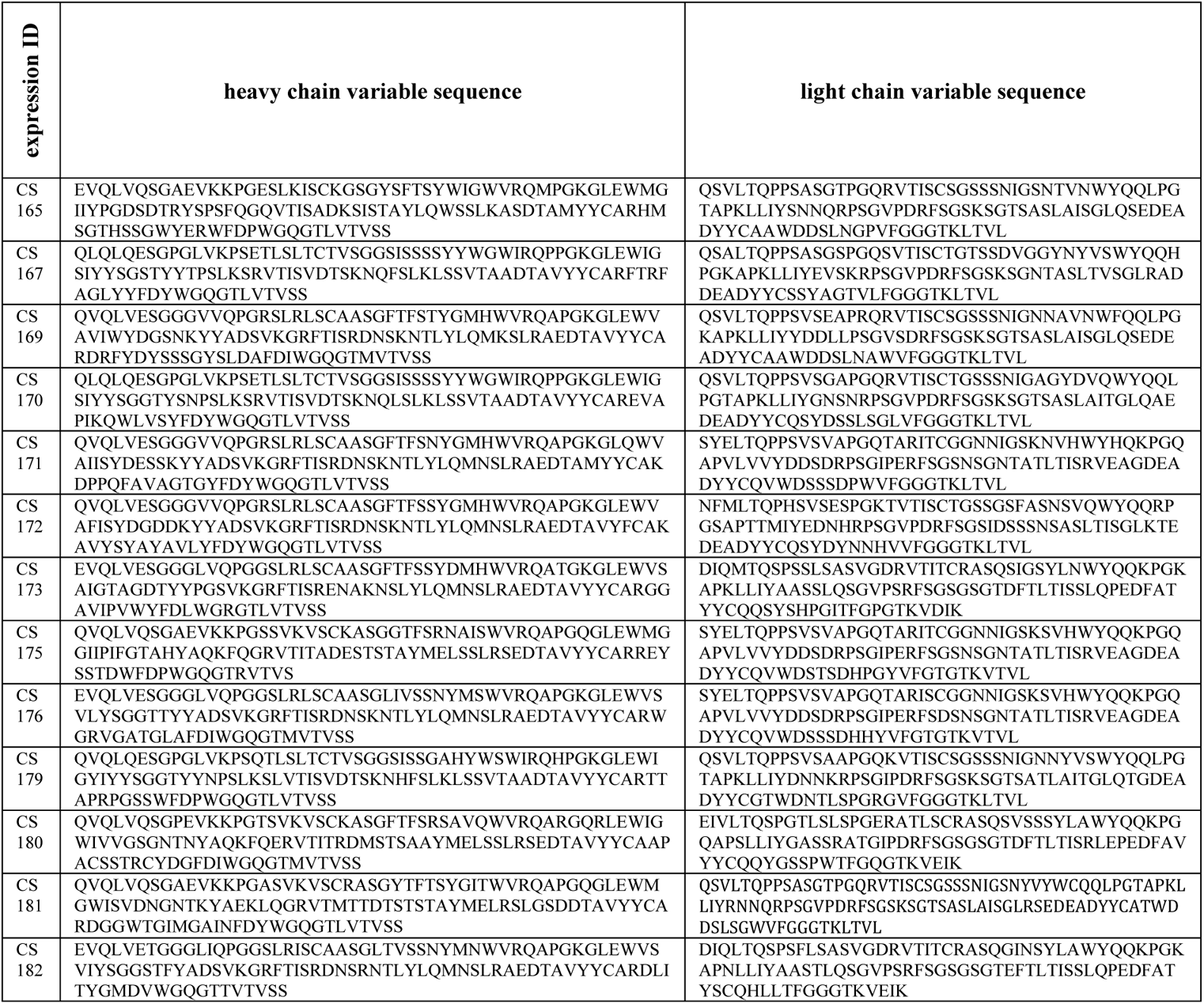
Sequences of antibodies reactive to RBD Beta in this study.

**Table S4.** X-ray data collection and refinement statistics.

| **Data collection** |  |  |  |
| --- | --- | --- | --- |
|  | CS23 + Beta RBD | CS44 + Beta RBD + COVA1-16 | CV07-287 + non-VOC RBD + COVA1-16 |
| Beamline | ALS 5.0.1 | SSRL 12-1 | SSRL 12-1 |
| Wavelength (Å) | 0.97741 | 0.97946 | 0.97946 |
| Space group | P 4<sub>1</sub> 2<sub>1</sub> 2 | P 2<sub>1</sub> 2<sub>1</sub> 2<sub>1</sub> | P 2<sub>1</sub> 2<sub>1</sub> 2<sub>1</sub> |
| Unit cell parameters |  |  |  |
| a, b, c (Å) | 145.9, 145.9, 112.0 | 89.6, 121.0, 138.0 | 89.6, 121.5, 136.9 |
| α, β, γ (°) | 90, 90, 90 | 90, 90, 90 | 90, 90, 90 |
| Resolution (Å)<sup>a</sup> | 50.0–2.86 (2.96–2.86) | 50.0–2.89 (2.99–2.89) | 50.0–2.45 (2.54–2.45) |
| Unique reflections<sup>a</sup> | 28,468 (2,764) | 34,085 (3,312) | 55,558 (5,448) |
| Redundancy<sup>a</sup> | 24.5 (14.8) | 10.3 (7.5) | 12.5 (6.6) |
| Completeness (%)<sup>a</sup> | 100.0 (99.5) | 98.9 (99.5) | 99.9 (99.2) |
| <I/σI><sup>a</sup> | 9.2 (1.0) | 14.0 (1.8) | 19.3 (1.1) |
| R<sub>sym</sub><sup>b</sup> (%)<sup>a</sup> | 25.6 (>100) | 16.9 (83.2) | 11.5 (>100) |
| R<sub>pim</sub><sup>b</sup> (%)<sup>a</sup> | 5.2 (35.2) | 5.3 (31.0) | 3.3 (48.0) |
| CC<sub>1/2</sub><sup>c</sup> (%)<sup>a</sup> | 98.2 (56.6) | 98.4 (78.6) | 97.4 (49.6) |
| **Refinement statistics** |  |  |  |
| Resolution (Å) | 46.9–2.86 | 40.6–2.89 | 40.5–2.45 |
| Reflections (work) | 28,466 | 34,078 | 55,547 |
| Reflections (test) | 2,000 | 1,687 | 2,728 |
| R<sub>cryst</sub><sup>d</sup> / R<sub>free</sub><sup>e</sup> (%) | 20.8/24.7 | 21.6/25.7 | 20.7/23.8 |
| No. of atoms | 4,783 | 8,323 | 8,512 |
| Macromolecules | 4,667 | 8,196 | 8,191 |
| Ligands | 87 | 89 | 96 |
| Solvent | 29 | 38 | 225 |
| Average B-value (Å<sup>2</sup>) | 53 | 61 | 60 |
| Macromolecules | 52 | 61 | 60 |
| Ligands | 80 | 63 | 64 |
| Solvent | 34 | 37 | 51 |
| Wilson B-value (Å<sup>2</sup>) | 49 | 57 | 51 |
| **RMSD from ideal geometry** |  |  |  |
| Bond length (Å) | 0.003 | 0.005 | 0.003 |
| Bond angle (°) | 0.54 | 0.90 | 0.66 |
| **Ramachandran statistics<sup>f</sup>** |  |  |  |
| Favored | 97.5 | 97.0 | 96.6 |
| Outliers | 0.00 | 0.19 | 0.00 |
| **PDB code** | **7S5P** | **7S5Q** | **7S5R** |

<sup>a</sup> Numbers in parentheses refer to the highest resolution shell.
<sup>b</sup> $R_{\text{sym}} = \sum_{hkl} \sum_i |I_{hkl,i} - \langle I_{hkl} \rangle| / \sum_{hkl} \sum_i I_{hkl,i}$ and $R_{\text{pim}} = \sum_{hkl} (1/(n-1))^{1/2} \sum_i |I_{hkl,i} - \langle I_{hkl} \rangle| / \sum_{hkl} \sum_i I_{hkl,i}$ , where $I_{hkl,i}$ is the scaled intensity of the $i^{\text{th}}$ measurement of reflection h, k, l, $\langle I_{hkl} \rangle$ is the average intensity for that reflection, and $n$ is the redundancy.
<sup>c</sup> CC<sub>1/2</sub> = Pearson correlation coefficient between two random half datasets.
<sup>d</sup> $R_{\text{cryst}} = \sum_{hkl} |F_o - F_c| / \sum_{hkl} |F_o| \times 100$ , where $F_o$ and $F_c$ are the observed and calculated structure factors, respectively.
<sup>e</sup> $R_{\text{free}}$ was calculated as for $R_{\text{cryst}}$ , but on a test set comprising 5% of the data excluded from refinement.
<sup>f</sup> From MolProbity (44).

**Table S5.** Hydrogen bonds and salt bridges identified at the antibody-RBD interface using the PISA program.

| CS23 | Dist.<br>[Å] | RBD |
| --- | --- | --- |
| **Hydrogen bonds** |  |  |
| H:THR28[N] | 3.1 | ALA475[O] |
| H:SER31[OG] | 3.3 | ALA475[O] |
| H:ASN32[ND2] | 3.2 | ALA475[O] |
| H:TYR33[OH] | 2.7 | LEU455[O] |
| H:SER53[N] | 3.3 | TYR421[OH] |
| H:SER53[OG] | 2.8 | ARG457[O] |
| H:SER53[OG] | 3.3 | LYS458[O] |
| H:SER53[OG] | 2.9 | TYR473[OH] |
| H:GLY54[N] | 2.9 | TYR421[OH] |
| H:SER56[OG] | 2.6 | ASP420[OD2] |
| H:TYR58[OH] | 3.5 | THR415[OG1] |
| H:ARG94[NH1] | 2.9 | ASN487[OD1] |
| H:TYR100[OH] | 3.9 | GLN493[OE1] |
| H:GLY26[O] | 2.7 | ASN487[ND2] |
| H:SER31[O] | 2.8 | TYR473[OH] |
| L:GLN31[NE2] | 3.2 | TYR505[OH] |
| L:SER93[N] | 3.5 | TYR505[OH] |
| L:SER93[OG] | 3.1 | ASP405[OD2] |
| L:ASP95A[OD1] | 2.9 | ARG408[NH2] |
| L:SER93[O] | 2.4 | ARG408[NH2] |
| L:TRP91[O] | 3.2 | TYR505[OH] |
| **Salt bridges** |  |  |
| L:ASP95A[OD2] | 3.9 | ARG408[NE] |
| L:ASP95A[OD1] | 2.9 | ARG408[NH2] |
| L:ASP95A[OD2] | 3.5 | ARG408[NH2] |

| CS44 | Dist.<br>[Å] | RBD |
| --- | --- | --- |
| **Hydrogen bonds** |  |  |
| H:CYS100B[N] | 2.7 | ALA475[O] |
| H:GLY53[O] | 3.9 | GLN493[NE2] |
| H:SER54[O] | 3.6 | GLN493[NE2] |
| H:GLY100[O] | 2.7 | TYR473[OH] |
| H:HIS100C[O] | 2.9 | ASN487[ND2] |
| H:ASP100D[OD1] | 3.0 | SER477[N] |
| H:ASP100D[OD1] | 3.4 | THR478[N] |
| H:ASP100D[OD1] | 3.3 | THR478[OG1] |
| H:ASP100D[OD2] | 2.5 | SER477[OG] |
| H:CYS100B[N] | 2.7 | ALA475[O] |
| H:GLY53[O] | 3.9 | GLN493[NE2] |
| H:SER54[O] | 3.6 | GLN493[NE2] |
| H:GLY100[O] | 2.7 | TYR473[OH] |
| H:HIS100C[O] | 2.9 | ASN487[ND2] |
| H:ASP100D[OD1] | 3.0 | SER477[N] |
| H:ASP100D[OD1] | 3.4 | THR478[N] |
| H:ASP100D[OD1] | 3.3 | THR478[OG1] |
| H:ASP100D[OD2] | 2.5 | SER477[OG] |
| H:CYS100B[N] | 2.7 | ALA475[O] |
| L:TYR32[OH] | 3.4 | PHE486[O] |

| CV07-287 | Dist.<br>[Å] | RBD |
| --- | --- | --- |
| **Hydrogen bonds** |  |  |
| H:CYS100B[N] | 3.0 | ALA475[O] |
| H:THR100[O] | 3.9 | TYR473[OH] |
| H:ASN100A[OD1] | 3.8 | SER477[N] |
| H:ASP100D[OD1] | 3.0 | SER477[N] |
| H:ASP100D[OD2] | 2.6 | SER477[OG] |
| H:ASP100D[OD1] | 3.5 | THR478[N] |
| H:ASP100D[OD1] | 3.3 | THR478[OG1] |
| H:TYR100C[O] | 2.9 | ASN487[ND2] |
| H:SER54[O] | 3.6 | GLN493[NE2] |
| H:CYS100B[N] | 3.0 | ALA475[O] |
| H:THR100[O] | 3.9 | TYR473[OH] |
| H:ASN100A[OD1] | 3.8 | SER477[N] |
| H:ASP100D[OD1] | 3.0 | SER477[N] |
| H:ASP100D[OD2] | 2.6 | SER477[OG] |
| H:ASP100D[OD1] | 3.5 | THR478[N] |
| H:ASP100D[OD1] | 3.3 | THR478[OG1] |
| H:TYR100C[O] | 2.9 | ASN487[ND2] |
| H:SER54[O] | 3.6 | GLN493[NE2] |
| H:CYS100B[N] | 3.0 | ALA475[O] |
| H:THR100[O] | 3.9 | TYR473[OH] |
| H:ASN100A[OD1] | 3.8 | SER477[N] |

